# PIF4-mediated regulation of H_2_O_2_ homeostasis controls Arabidopsis seedling thermomorphogenesis

**DOI:** 10.1101/2025.09.02.673620

**Authors:** Ritesh Kumar Raipuria, Kumud Saini, Mahesh Kumar Panda, Aditi Dwivedi, Aashish Ranjan

## Abstract

Thermomorphogenesis under high ambient temperature involves extensive developmental changes, including hypocotyl elongation at the seedling stage, in Arabidopsis. Reactive Oxygen Species (ROS), particularly hydrogen peroxide (H_2_O_2_), are important signaling molecules, playing crucial roles in plant development and stress responses. While ROS-homeostasis is shown to be crucial for maintaining cellular functions and mediating various developmental responses, the involvement of ROS-homeostasis in the regulation of thermomorphogenic responses and the underlying genetic basis remains poorly understood. In this study, comprehensive transcriptomic analyses revealed strong induction of ROS homeostasis and signaling genes in Arabidopsis seedlings under high ambient temperature. Pharmacological and genetic experiments showed that maintaining H_2_O_2_ homeostasis is crucial for seedling thermomorphogenesis. We identified PHYTOCHROME INTERACTING FACTOR 4 (PIF4) as a key regulator of H_2_O_2_ homeostasis via direct transcriptional activation of *CAT2* and *CAT3* genes, which are involved in the regulation of H_2_O_2_ levels, to modulate hypocotyl elongation under high temperature. Genetic and biochemical experiments confirmed that CATs act downstream to PIF4 in the same signaling pathway to regulate high-temperature-responsive hypocotyl elongation. Interestingly, elevated H_2_O_2_ levels reduced PIF4 protein abundance under high temperature. Together, our findings establish a PIF4-CAT-H_2_O_2_ regulatory module, functioning alongside the canonical PIF4-Auxin module, that integrates to auxin signaling to fine-tune hypocotyl elongation under high temperature by maintaining H_2_O_2_ homeostasis.

**Highlights:**

- High ambient temperature significantly affects ROS homeostasis and signaling genes in Arabidopsis seedlings.
- H_2_O_2_ homeostasis is crucial for high-temperature-mediated hypocotyl elongation, a signature feature of Arabidopsis thermomorphogenesis.
- PHYTOCHROME INTERACTING FACTOR 4 (PIF4) regulates the expression of *CATALASE* genes in a temperature-dependent manner to maintain H_2_O_2_ levels for seedling thermomorphogenesis.
- Elevated H_2_O_2_ level reduces PIF4 protein abundance, thus forming a PIF4-CAT-H_2_O_2_ regulatory module that integrates with auxin to fine-tune hypocotyl elongation under high temperature.

## Introduction

Temperature is one of the most important environmental factors for plant growth and development. While extreme heat stress is detrimental for plants due to denaturation and folding defects of key enzymes and proteins, a slight (5-6 °C) increase in ambient temperature significantly affects plant development. Plants exhibit remarkable developmental plasticity, as a series of morphological adaptive responses, to high ambient temperatures.^1–3^ These adaptive responses, which are collectively called thermomorphogenesis, include hypocotyl and petiole elongation, reduced leaf area, and accelerated flowering in the model plant Arabidopsis. Hypocotyl elongation is a signature thermomorphogenic response at the seedling stage that is suggested to protect the shoot apical meristem (SAM) and emerging leaves from ambient soil warmth. ^4^

Arabidopsis hypocotyl elongation in response to high ambient temperature involves thermoreceptors and downstream signalling components, including transcription factors and phytohormones. High temperature is reported to be sensed by active Pfr form of Phytochrome B (PHYB), EARLY FLOWERING 3 (ELF3), PIF7 mRNA, and ARABIDOPSIS FUS3-COMPLEMENTING (AFC) kinase. ^5–9^ In addition, THERMO-WITH ABA-RESPONSE 1 (TWA1), a transcriptional co-regulator, was recently discovered as a thermosensor under heat stress. ^10^ Downstream to PHYB, PHYTOCHROME INTERACTING FACTORs (PIFs), basic-helix-loop-helix (bHLH) transcriptional factors, bind the promoters of genes positively regulating thermomorphogenesis. Among the eight PIFs in Arabidopsis, PIF4, PIF5, and PIF7 are shown to have roles in high-temperature-responsive hypocotyl elongation. ^11–15^ As a central regulator of thermomorphogenesis, PIF4 induces the expression of auxin biosynthetic genes, encoding flavin monooxygenase-like enzyme YUCCA8 (YUC8), TRYPTOPHAN AMINOTRANSFERASE OF ARABIDOPSIS (TAA1), and cytochrome-p450 family member CYP79B2, thereby increasing auxin levels. ^16–17^ Auxin, together with brassinosteroid, enhances cellular elongation, leading to hypocotyl elongation. ^18–20^ PIF7 is reported to regulate daytime high-temperature response in the early seedling growth, suggesting specific roles of different PIFs in Arabidopsis thermomorphogenesis. ^8^ While the PIF-mediated induction of auxin levels leading to hypocotyl elongation is extensively studied, crucial functions of other signalling molecules, such as sucrose and Reactive Oxygen Species (ROS), are envisaged for hypocotyl elongation in response to high temperature. ^21–22^

Reactive Oxygen Species (ROS), which include superoxide anion (O_2_^•−^), hydrogen peroxide (H_2_O_2_), singlet oxygen (^1^O_2_), and hydroxyl radical (OH^•^), are produced as by-products of several redox reactions during plant metabolism and accumulate under various stress responses. ^23–25^ ROS, in addition to their involvement in stress responses, are shown to function as important signaling molecules regulating plant growth and development. ^23, 26–29^ For example, the spatial pattern of ROS, particularly H_2_O_2_, regulates stomatal development as well as light-induced stomatal opening. ^30–31^ ROS homeostasis has been demonstrated to regulate cambial cell proliferation and tapetal cell differentiation of the anther in Arabidopsis. ^32–33^ Seed germination is positively correlated with increased ROS production, which is essential to break seed dormancy. ^34–35^ Since ROS are short-lived, highly reactive, and are capable of inducing programmed cell death, their levels are kept under tight control by balancing ROS production, transport, and scavenging to prevent unintended cellular oxidation. ^23, 28^ While Respiratory Burst Oxidase Homologs (RBOHs) are plasma membrane-localized enzymes involved in ROS production, several enzymes, such as superoxide dismutase (SOD), ascorbate peroxidase (APX), guaiacol peroxidase (GPX), and catalase (CAT), are involved in ROS scavenging. Catalases convert the H_2_O_2_ into water and oxygen molecules. ^36–38^ These ROS scavenging enzymes maintain ROS homeostasis under biotic and abiotic stresses by preventing excess ROS accumulation. ^39^

ROS is known to be extensively involved in plant responses to environmental fluctuations, such as changes in light and temperature. Cross-talk between light and ROS signaling has been well established. For example, Phytochrome B (PHYB), a light and temperature sensor, promotes rapid ROS accumulation under high light and other abiotic stresses. ^40^ Interestingly, a report suggests the involvement of H_2_O_2_ in seedling establishment under dark conditions. ^41^ While PIF1 and PIF3 hinder ROS accumulation, the WRKY33-PIF4 regulatory loop is essential to maintain H_2_O_2_ homeostasis for seedling growth and development. ^42–43^ Arabidopsis seedlings exhibit increased accumulation of O_2_^•−^ and H_2_O_2_ in the hypocotyl and decreased O_2_^•−^ levels in the cotyledons upon shade treatment. ^44^ Heat stress has been shown to elevate ROS levels and activate stress signaling pathways. ^45–46^ The accumulation of H_2_O_2_ induces the expression of various heat shock factors (HSFs), heat shock proteins (HSPs), and ROS scavenging enzymes that help plants to survive under heat stress. ^47–49^ ROS burst induced by heat stress slows down the shoot apical meristem (SAM) maturation, thereby delaying the transition of the vegetative stage to achieve heat-stress resilience. ^50^ Among the various antioxidant enzymes, CATALASE 2 (CAT2) plays a key role in short and long-term heat tolerance. ^51^ A recent report suggests that glutathione, another potent antioxidant maintaining cellular redox state, plays a crucial role in thermomorphogenesis and heat stress tolerance. ^22^

While the role of ROS and the antioxidant system in mitigating heat stress has been extensively studied, the information on the involvement of ROS in thermomorphogenesis is rather limited. In this study, we systematically investigated the involvement of ROS homeostasis and underlying genetic regulation for developmental plasticity in thermomorphogenesis. By using pharmacological, molecular, and genetic approaches, we show that H_2_O_2_ homeostasis is essential for high-temperature-mediated hypocotyl elongation. We identified PIF4 as a key genetic regulator for maintaining H_2_O_2_ levels under high temperature via transcription regulation of *CATALASE* genes. Together, our study integrates high ambient temperature and ROS signaling for developmental plasticity under high ambient temperatures.

## Results

### High ambient temperature induces significant changes in ROS homeostasis and signaling

To investigate the global effects of high ambient temperature on ROS homeostasis and signaling, we analyzed multiple publicly available transcriptomic datasets that investigated the effects of high temperature on Arabidopsis seedlings. ^15, 22, 52^ As expected, gene ontology enrichment analyses of upregulated genes across these datasets consistently identified the enrichment of categories such as “response to temperature stimulus” and “response to auxin-activated signaling pathway”. Interestingly, categories such as “response to reactive oxygen species”, “response to hydrogen peroxide”, and “response to oxygen-containing compound” were also significantly enriched among the upregulated genes at different time points (Figure 1A). Furthermore, high temperature treatment led to differential expression of key ROS scavenging genes, such as *CAT1* and *CAT3*, among the DEGs. ^22^ Together, the meta-analysis of three independent transcriptomic studies suggested an involvement of ROS homeostasis and signaling in regulating Arabidopsis seedling thermomorphogenesis.

**Figure 1.**
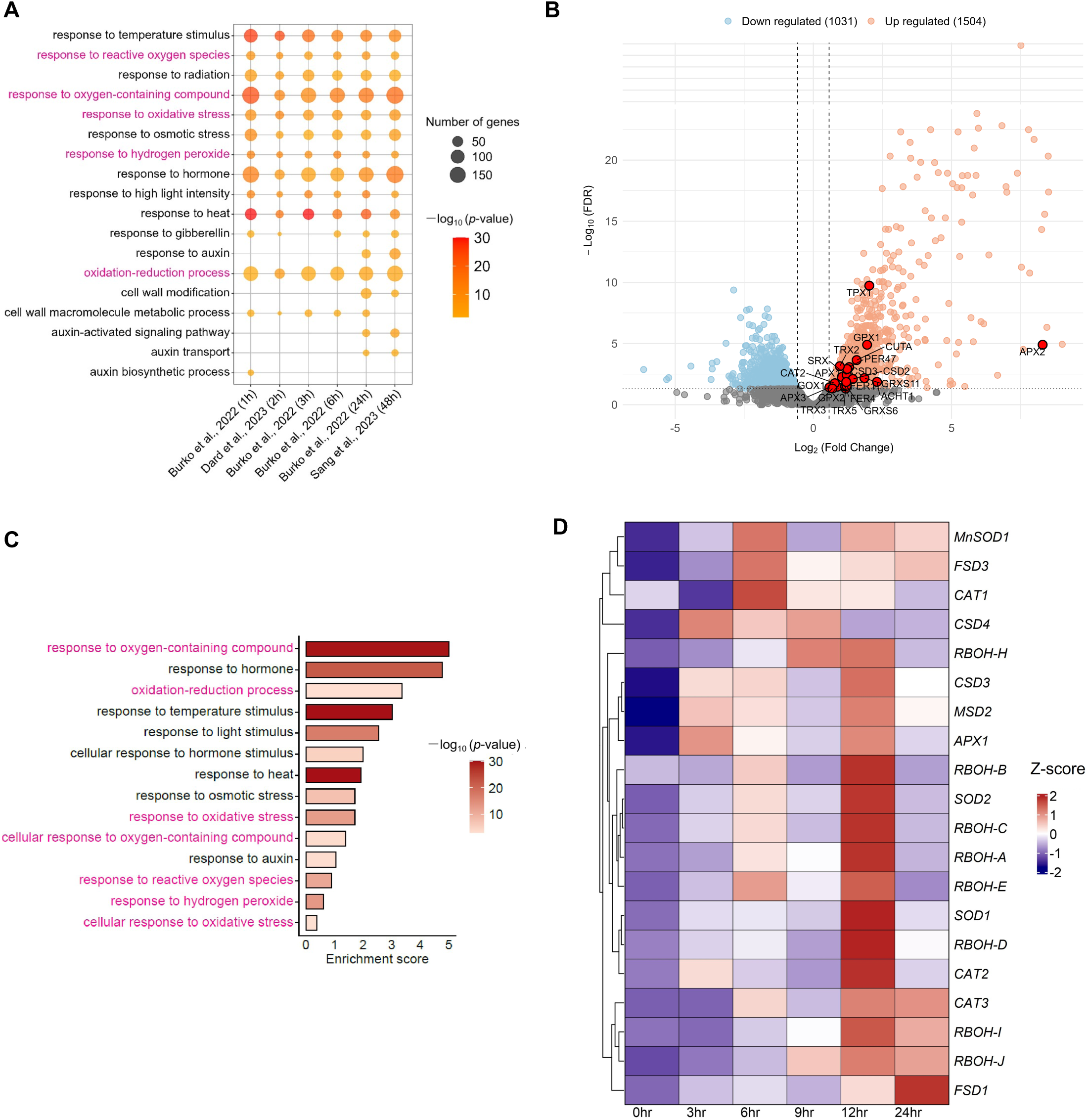
High ambient temperature significantly induces the expression of ROS homeostasis and signaling genes. (A) Enriched biological processes for the genes upregulated in response to high temperature at different time points. The Gene Ontology (GO) enrichment analysis with a significance threshold of *P*<0.005 was performed for upregulated genes from three independent studies: Burko et al., 2022 (1hr, 3hrs, 6hrs, 24hrs), Dard et al., 2023 (2hrs), and Sang et al., 2023 (48hrs). The dot size represents the number of genes upregulated in each GO category, and the color gradient reflects the statistical significance as –log10 (*P*-value). GO terms highlighted in magenta represent processes related to ROS homeostasis and signaling. (B) Volcano plot showing differentially expressed genes (DEGs) in hypocotyl after 1hour high temperature treatment. Genes with significant upregulation (FDR < 0.05 and log2FC > 0.585) are shown in salmon pink color, while significantly downregulated genes (FDR < 0.05 and log2FC < 0.585) are shown in sky blue color. Selected ROS homeostasis and signaling genes are highlighted as red bold circles and labeled. (C) Enriched biological processes (*P*<0.005) for the genes upregulated in hypocotyl after 1hour high temperature treatment. The size of the bars represents the number of genes upregulated in each GO category, and the color gradient reflects the statistical significance as –log10 (*P*-value). GO terms highlighted in magenta represent ROS-related processes. (D) Expression patterns of ROS-homeostasis genes in Arabidopsis hypocotyl at different time points in response to high ambient temperature. The heatmap represents Z-score transformed values of fold change (2^-ΔΔCT^) of each gene under different time points compared to the control condition.

Since hypocotyl elongation is the signature feature of seedling thermomorphogenesis, we performed RNA-seq analysis to investigate the effects of high ambient temperature on ROS homeostasis and signaling genes specifically in Arabidopsis hypocotyl. High temperature treatment of 1 hour caused differential expression of 2529 genes in hypocotyl, with upregulation of 1500 genes and downregulation of 1029 genes (Figure 1B, Supplemental Dataset S1). Gene ontology enrichment analysis of upregulated genes showed enrichment of categories such as “response to heat”, “response to light stimulus”, “response to oxidative stress”, “response to hydrogen peroxide”, and “response to reactive oxygen species”, in addition to categories related to temperature, hormones, and growth (Figure 1C, Supplemental Dataset S2). Genes encoding enzymes involved in H_2_O_2_ production and scavenging, such as *CSOD2*, *CSOD3*, and *CAT2*, were induced in hypocotyl under high temperature (Figure 1B, Supplemental Dataset S1). These results further strengthened the role of ROS, particularly H_2_O_2_, in regulating high-temperature-responsive hypocotyl elongation.

Since RBOHs, SODs, and CATs are the key enzymes responsible for ROS homeostasis, we then performed the expression analysis of *RBOH*s, *SOD*s, and *CAT*s in response to high temperature at temporal resolution in the hypocotyl. Eight out of ten *RBOH*s and all but one *SOD*s showed significant up-regulation at 12 hours after high temperature treatment compared to seedlings at control temperature (Figure 1D). The three *CAT* genes were also significantly upregulated at 12 hours post high temperature treatment. Together, these expression profiles showed the significant effect of high ambient temperatures on ROS homeostasis genes and highlight 12 hours as an important time-point for molecular changes associated with ROS levels and homeostasis in response to high temperature.

### Hydrogen peroxide (H_2_O_2_) is an important regulator of Arabidopsis seedling thermomorphogenesis

Since H_2_O_2_ is the most stable ROS and is known to regulate various developmental and stress responses, we investigated the involvement of H_2_O_2_ in temperature-mediated hypocotyl elongation. DAB (3,3’-Diaminobenzidine) staining showed significantly higher H_2_O_2_ accumulation in seedlings grown at 28 °C as compared to the control seedlings grown at 22°C, suggesting a potential involvement of H_2_O_2_ in hypocotyl elongation under high temperature (Figures 2A–2B). We then performed pharmacological experiments using N, N′- Dimethylthiourea (DMTU), a ROS scavenger that would reduce H_2_O_2_ levels, to test the requirement of H_2_O_2_ for high-temperature-mediated hypocotyl elongation. High temperature-mediated hypocotyl elongation was attenuated with increasing DMTU concentrations in a dose-dependent manner, confirming the requirement of H_2_O_2_ for the elongation response (Figure 2C). Mock seedlings under control temperature, however, did not show a significant change in hypocotyl length in response to exogenous DMTU treatment. We further performed exogenous treatment experiments using 3-Amino-1,2,4-triazole (3AT), a CATALASE inhibitor that would increase H_2_O_2_ levels. Lower concentrations of 3AT (0.05 and 0.1 µM) induced temperature-mediated hypocotyl elongation (Figure 2D). However, the hypocotyl elongation under high temperature subsided with increasing 3AT concentration beyond 0.1 µM. These results showed that the increasing H_2_O_2_ levels promote hypocotyl elongation up to a range, with attenuation of temperature-induced elongation at H_2_O_2_ levels beyond a threshold. Similar to DMTU treatment, 3AT at different concentrations did not show any significant change in hypocotyl length under control temperature (Figures 2C–2D). We quantified the H_2_O_2_ levels in 3AT- and DMTU-treated seedlings to confirm the desired manipulation by the exogenous treatments. As expected, we observed higher H_2_O_2_ accumulation in 3AT-treated seedlings and lower H_2_O_2_ accumulation in DMTU-treated seedlings in comparison to untreated seedlings (Figure S1). Taken together, these results showed that a proper H_2_O_2_ homeostasis, not very high or low H_2_O_2_ levels, is required for high-temperature-responsive Arabidopsis hypocotyl elongation.

**Figure 2.**
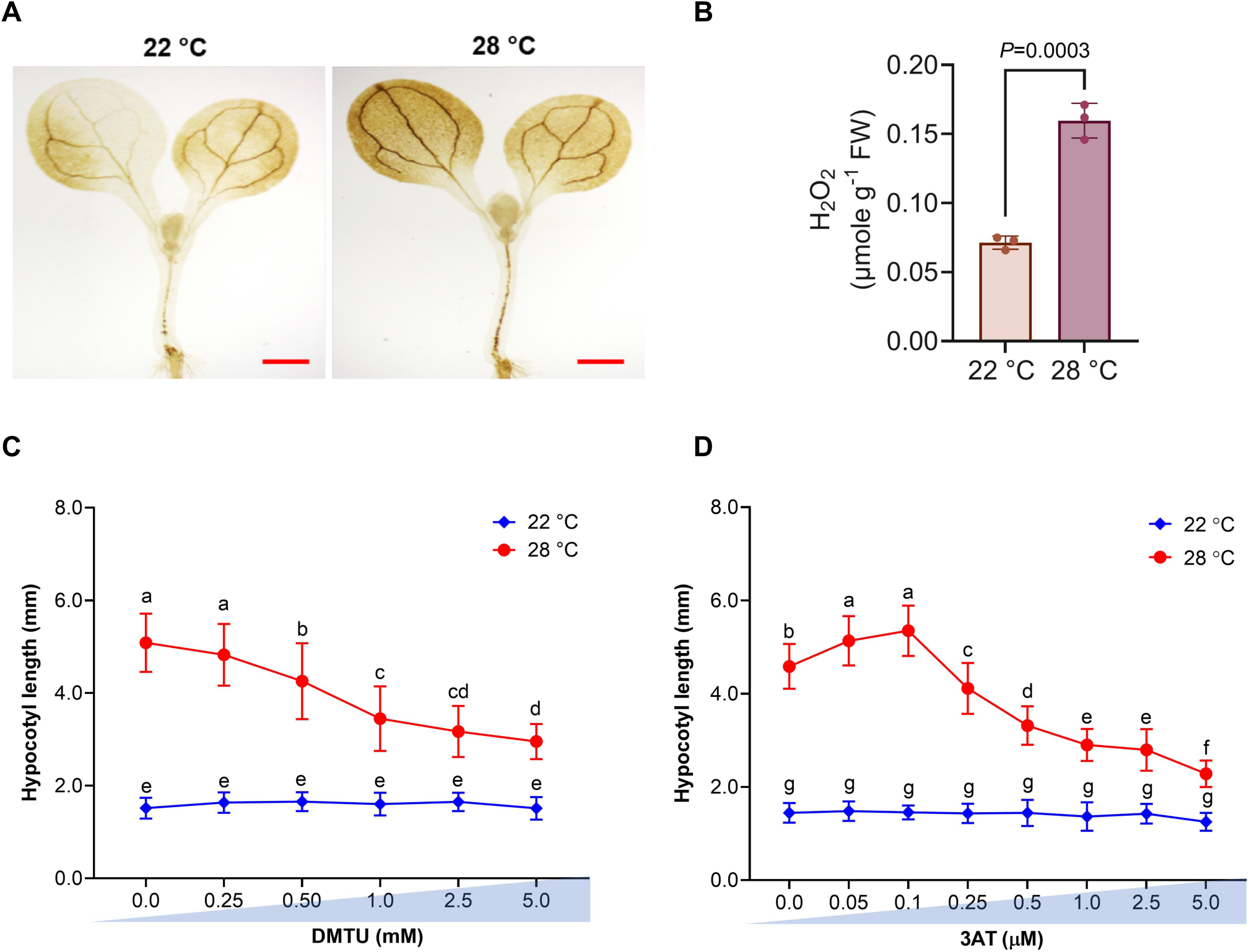
H_2_O_2_ homeostasis is essential for high-temperature-responsive hypocotyl elongation. (A) DAB staining showing H_2_O_2_ accumulation in wild-type Arabidopsis seedlings grown under control (22 °C) and high temperature (28 °C for 12hrs) conditions. Scale bar-1 mm. (B) Quantification of H_2_O_2_ levels in wild-type seedlings grown under control and high temperature conditions. Error bars represent standard deviation (SD). Statistical significance between the two samples was calculated using a two-tailed Student’s t-test. (C) DMTU dosage response curve for hypocotyl elongation under control and high temperature conditions. (D) 3AT-dosage response curve for hypocotyl length under control and high temperature conditions. Wild-type seedlings were grown in control conditions for 4 days and then transferred to the media plates with different concentrations of 3AT (0.05 to 5.0 µM) or DMTU (0.25 to 5.0 mM), followed by quantification of hypocotyl length after 72hrs. Error bars represent SD. Different lowercase letters above the error bars represent statistically significant differences among the samples as determined by two-way ANOVA followed by Tukey’s post hoc test, *P*<0.01.

### Catalase-mediated control of H_2_O_2_ levels modulates thermomorphogenic response

ROS scavenging is crucial for maintaining proper ROS homeostasis for optimal plant developmental and stress responses. ^29, 35^ Since catalases (CATs) are one of the most important H_2_O_2_ scavenging enzymes, we checked the involvement of genes encoding CATs in high-temperature-mediated hypocotyl elongation. Among the three genes encoding CATs in Arabidopsis, *CAT1*, *CAT2,* and *CAT3*, we primarily focused on *CAT2* and *CAT3*, as both of them showed maximal induction in gene expression at 12 hrs post high temperature treatment along with several *RBOH*s and *SOD*s that are important for ROS-homeostasis (Figure 1C). Moreover, *CAT1* is primarily expressed in seeds and pollen. ^53–54^ We evaluated the hypocotyl elongation phenotype of mutants as well as overexpression lines of *CAT2* and *CAT3,* along with wild-type (Col-0). Two independent mutants of the *CAT2* gene, *cat2-1* and *cat2-2*, exhibited attenuated hypocotyl elongation under high temperature in comparison with Col-0 (Figures 3A and 3C). Interestingly, three independent *35S:CAT2* overexpression lines also showed attenuated hypocotyl elongation under high temperature. Similarly, the *cat3* mutant and three independent *35S:CAT3* overexpression lines also showed attenuated elongation response compared to Col-0 under high temperature (Figures 3B and 3D). To connect the mutant and overexpression phenotypes to H_2_O_2_, we quantified H_2_O_2_ accumulation in these lines. As expected, *cat2* and *cat3* mutants showed higher H_2_O_2_ accumulation, and overexpression lines of CAT2 and CAT3 had reduced H_2_O_2_ levels compared to Col-0 under high temperature (Figure S2). Interestingly, *cat2* mutants accumulated higher levels of H_2_O_2_ than *cat3*, suggesting stronger effects of *cat2* mutation than *cat3*. To further investigate the genetic interaction of CAT2 and CAT3, we generated a *cat2-1cat3* double mutant that exhibited attenuated hypocotyl elongation response, similar to *cat2* mutants (Figures 3A–3D). Moreover, *cat2-1* and *cat2-2* mutants showed stronger attenuation response than the *cat3* mutant. These suggest that CAT2 may play a prominent role over CAT3 for hypocotyl elongation response under high temperature. Overall, these genetic results show the important roles of *CAT2* and *CAT3* in temperature-mediated hypocotyl elongation. Furthermore, quantification of H_2_O_2_ in mutant and overexpression lines and associated hypocotyl phenotype supported the importance of H_2_O_2_ homeostasis for seedling thermomorphogenesis.

**Figure 3.**
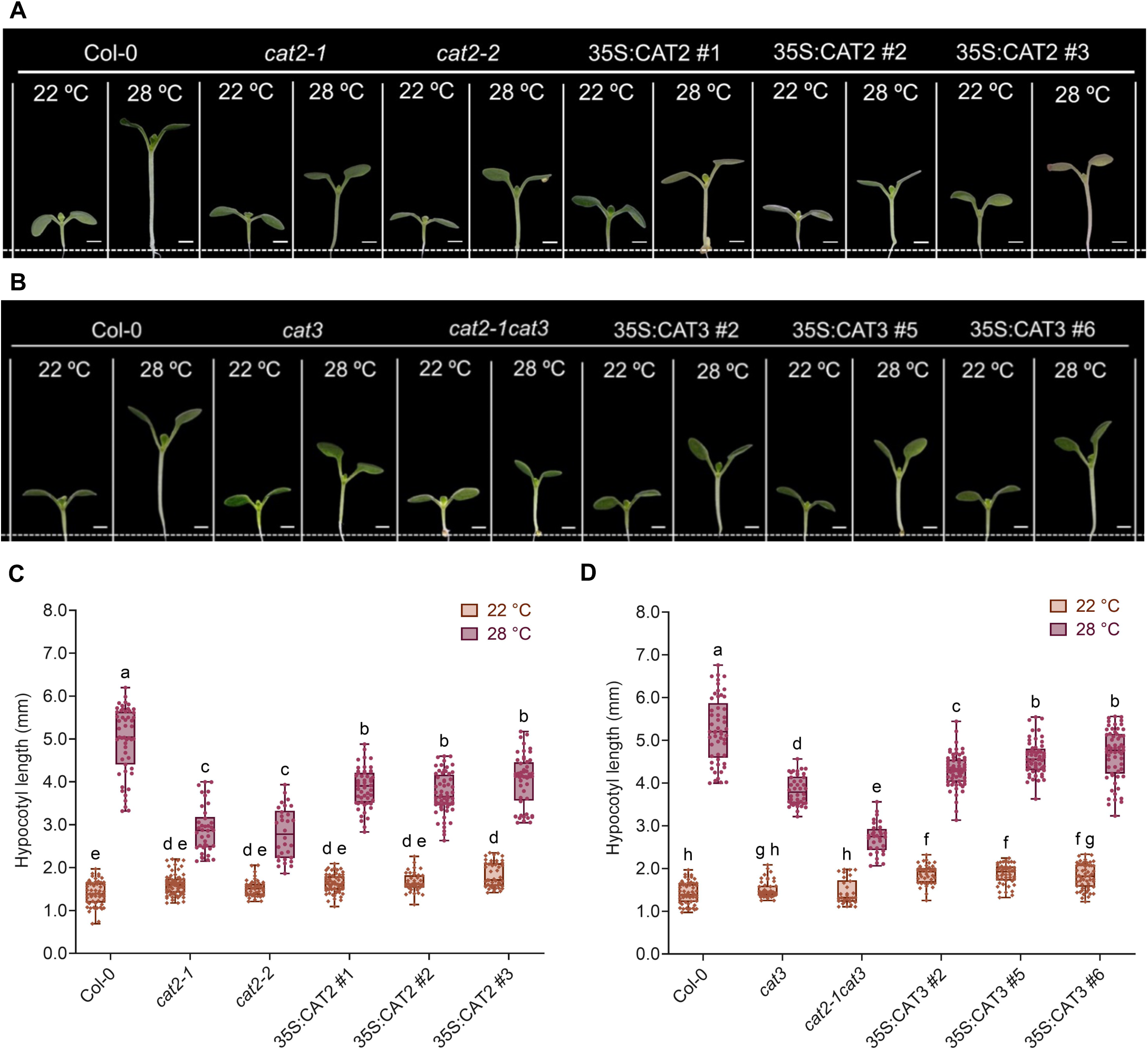
Genetic manipulation of the *CAT2* and *CAT3* genes impairs temperature-mediated hypocotyl elongation. (A and B) Representative seedling images showing hypocotyl elongation of Col-0, *cat2* and *cat3* mutants, *cat2cat3* double mutant, and *CAT2* and *CAT3* over-expression lines grown under control (22 °C) and high temperature (28 °C) conditions. Scale bar-1mm. (C and D) Box plot showing hypocotyl length quantification of Col-0, *cat* mutants, and CAT over-expression lines grown under control and high temperature conditions. Error bars represent SD. Central horizontal lines within each box represent the mean. Small circles and diamonds on the box represent the number of seedlings quantified. Different lowercase letters above the bars indicate statistically significant differences among the samples, as determined by two-way ANOVA followed by Tukey’s post hoc test (*P*<0.01).

### PIF4 induces *CAT2* and *CAT3* transcription under high ambient temperature

High temperature-mediated induction of *CAT2* and *CAT3* expression and attenuated hypocotyl elongation of *cat2, cat3,* and *cat2-1cat3* mutants suggested the possibility of transcriptional regulation of *CAT* genes under high temperature. Therefore, we scanned the promoters of *CAT2* and *CAT3* for potential transcription factor (TF) binding sites and detected three G-boxes each in the promoters of *CAT2* and *CAT3* genes (Figure S3). Since PIF4, a master transcriptional regulator of Arabidopsis thermomorphogenesis, binds to the G-box in the promoters of the target genes ^8, 11, 13^, we investigated the possibility of PIF4-mediated regulation of *CAT2* and *CAT3* under high temperature. While high temperature significantly induced the *CAT2* and *CAT3* expression in Col-0, the two genes failed to show induction in *pif4* and *pif457* seedlings under high temperature, suggesting that the high temperature-mediated induction of *CAT2* and *CAT3* requires the presence of *PIF4* (Figure 4A). Moreover, a similar expression pattern of *CAT2* and *CAT3* in both *pif4* and *pif457* mutants highlights the major role of PIF4 in temperature-mediated induction of *CAT2* and *CAT3*. To further confirm the PIF4-dependent transcriptional regulation of *CAT2* and *CAT3* genes, we generated *pif4*/*proCAT2*:GUS and *pif4*/*proCAT3*:GUS lines that showed reduced GUS activity in cotyledons and hypocotyl under high temperature compared to the *proCAT2*:GUS and *proCAT3*:GUS lines, respectively (Figure 4B). To examine whether PIF4 directly regulates the transcription of *CAT2* and *CAT3* genes, we performed luciferase trans-activation and dual luciferase assays. Both assays showed that PIF4 activates the promoters of the *CAT2* and *CAT3* genes (Figures 4C–4D). To further confirm the direct binding of PIF4 to the G-boxes present in the promoters of the *CAT* gene, we performed Chromatin immunoprecipitation (ChIP)-PCR. As controls, we included the *IAA29* promoter (Known PIF4 targets) as a positive control and the *UBC30* promoter as a negative control. We got the enrichment of PIF4 from G-boxes of *CAT2* and *CAT3* promoters using anti-PIF4 antibody under high temperature (Figures 4E–4F). These results, together, confirm PIF4-mediated direct transcriptional regulation of *CAT2* and *CAT3* genes under high temperature.

**Figure 4.**
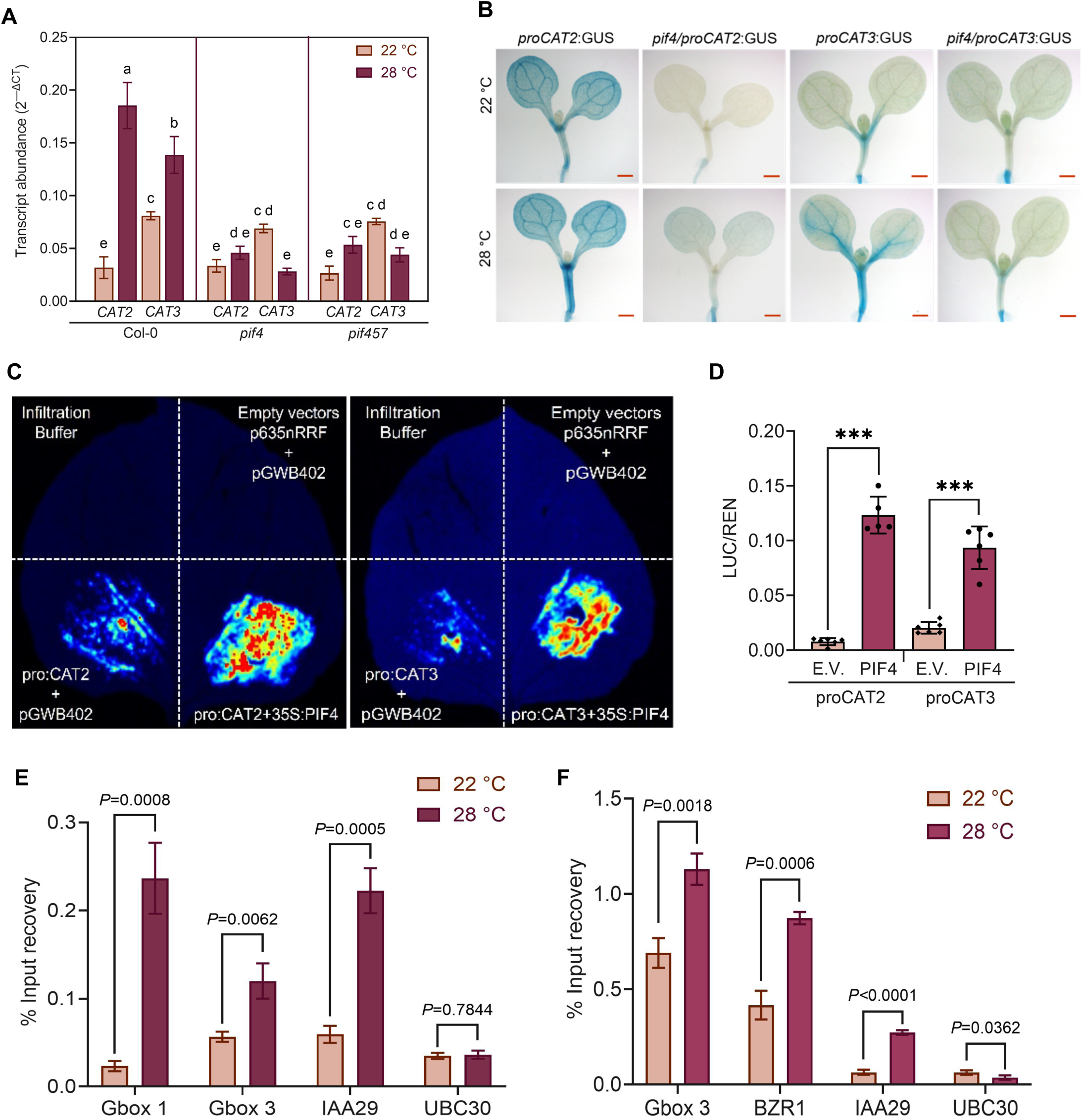
PIF4-mediated direct transcriptional regulation of *CAT2* and *CAT3* genes. (A) Transcript abundance (2^-ΔCT^) of *CAT2* and *CAT3* genes relative to endogenous control *GAPDH* in the seedlings of Col-0, *pif4,* and *pif457* mutants grown under control (22 °C) and high temperature (28 °C) conditions. Error bar represents SD (n=6). Different lowercase letters indicate statistically significant differences among the samples, as determined by two-way ANOVA followed by Tukey’s post hoc test (*P*<0.01). (B) Representative GUS-staining images of pro*CAT2*:GUS, *pif4*/pro*CAT2*:GUS, pro*CAT3*:GUS, and *pif4*/pro*CAT3*:GUS seedlings grown under control and high temperature conditions. Scale bar-100 µm, (C) Luciferase trans-activation assay showing activation of *CAT2* and *CAT3* promoters by PIF4 transcription factor in *Nicotiana benthamiana* leaves. (D) Bar plot showing LUC/REN ratio for the interaction of PIF4 with the promoters of *CAT2* and *CAT3* genes, where REN is used as an internal control. Error bars represent the SD (n=6). Asterisks above the bar indicate statistical significance between the samples using a two-tailed Student’s t-test (\*\*\**P*˂0.001). (E) ChIP-qPCR assay showing binding and enrichment of PIF4 from G-box 1 (– 1062 position from ATG) and G-box 3 (–2978 position from ATG) of the *CAT2* promoter under control and high temperature conditions. (F) ChIP-qPCR assay showing binding and enrichment of PIF4 from G-box 3 (–5236 position from ATG) of the *CAT3* promoter under control and high temperature conditions. *IAA29* and *BZR1* promoters were used as positive controls, whereas the *UBC* promoter was used as a negative control. Statistical significance between the samples, indicated above the bars, was calculated using a two-tailed Student’s t-test.

### Genetic interaction between PIF4 and CATs to regulate temperature-mediated hypocotyl elongation

Since PIF4 directly regulates transcription of *CAT2* and *CAT3*, we then quantified H_2_O_2_ levels in *pif4* and *pif457* mutants to investigate PIF4-mediated regulation of H_2_O_2_ homeostasis for hypocotyl elongation in high temperature. As expected, *pif4* and *pif457* seedlings show significantly higher H_2_O_2_ accumulation in both control and high temperature when compared with Col-0 seedlings (Figures 5A–5B). We hypothesized that elevated H_2_O_2_ level, beyond the threshold, in the *pif4* mutant could contribute to its failure to elongate hypocotyl under high temperature. To test this hypothesis, we treated the *pif4* mutant with DMTU, a ROS scavenger that would reduce H_2_O_2_ levels. Exogenous DMTU treatment resulted in a significant increase in the *pif4* hypocotyl elongation under high temperature (Figure S4). However, Col-0 seedlings showed a significant reduction in hypocotyl elongation under high temperature in response to DMTU treatment (Figure S4). These results further suggest that H_2_O_2_-homeostasis is critical for high-temperature-mediated hypocotyl elongation and that PIF4 is a key regulator of H_2_O_2_-homeostasis via transcriptional regulation of *CAT2* and *CAT3*.

**Figure 5.**
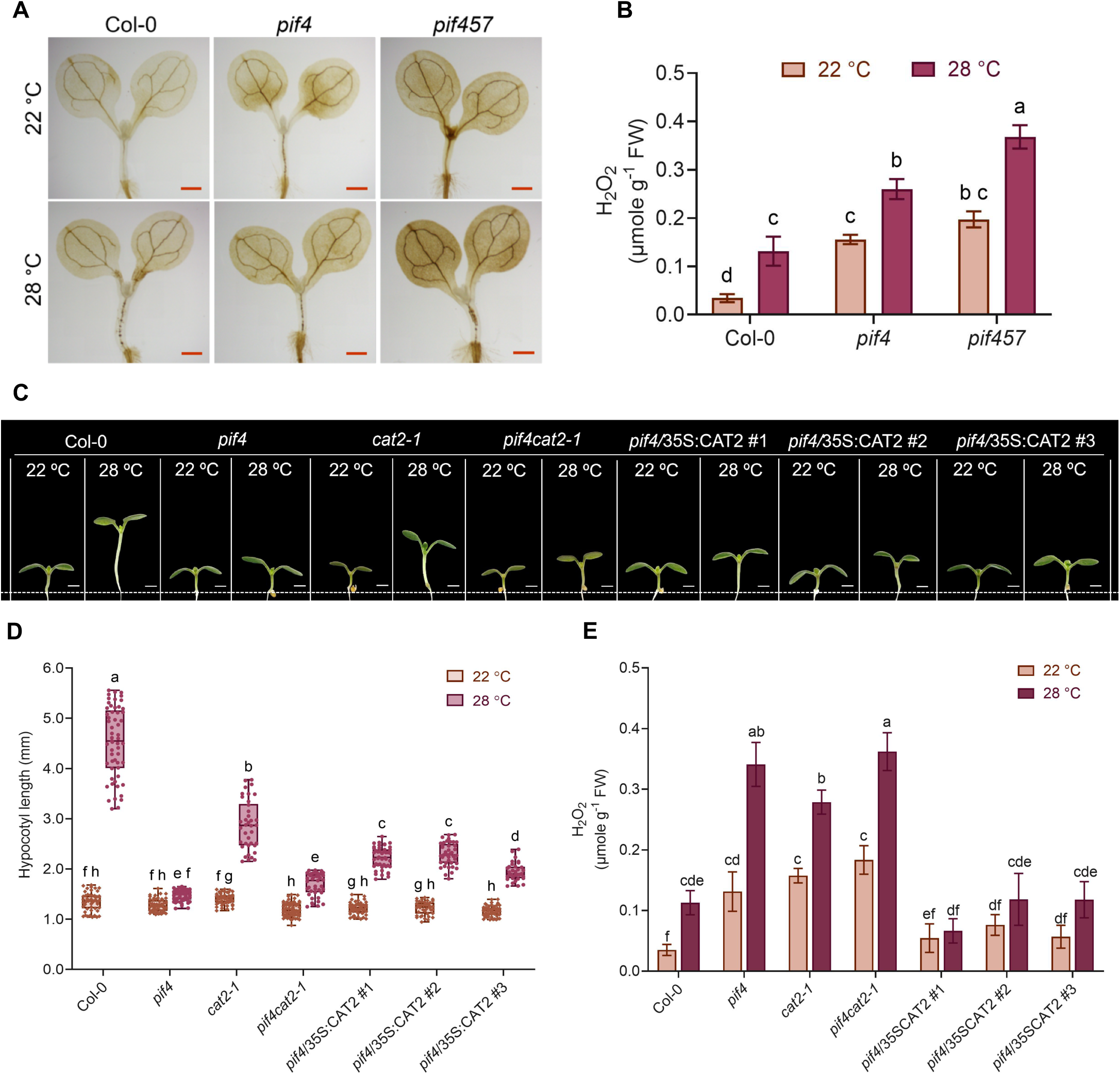
*PIF4* and *CAT2* genetically interact to control H_2_O_2_ levels and seedling thermomorphogenesis. (A) Representative images of Col-0, *pif4,* and *pif457* seedlings showing DAB staining under control (22 °C) and high temperature (28 °C) conditions. Scale bar-1 mm. (B) Quantification of H_2_O_2_ levels in Col-0, *pif4*, and *pif457* seedlings grown under control and high temperature conditions. (C) Representative seedling images showing hypocotyl elongation of Col-0, *pif4*, *cat2-1*, and *pif4cat2-1* double mutant, along with three independent *pif4*/35S:CAT2 lines grown under control and high temperature conditions. Scale bar - 1 mm. (D) Quantification of hypocotyl length of the genotypes shown in (C). (E) Quantification of H_2_O_2_ levels in the genotypes shown in (C) in control and high temperature conditions. Error bars in (B), (D), and (E) represent the SD from the mean values. Different lowercase letters indicate statistically significant differences among the samples, as determined by two-way ANOVA followed by Tukey’s post hoc test (*P*<0.01).

Next, we generated various genetic lines, such as *pif4cat2-1*, *pif4cat3*, *pif4*/35S:CAT2, and *pif4*/35S:CAT3, to investigate the genetic interaction between PIF4 and CATs to regulate H_2_O_2_ levels and hypocotyl elongation under high temperature. *pif4cat2-1* and *pif4cat3* double mutants showed a significant reduction in hypocotyl elongation when compared with mutants *cat2* and *cat3*, respectively (Figures 5C and 5D, S5A and S5B). Consistent with this, these *pif4cat2-1* and *pif4cat3* double mutants accumulated more H_2_O_2_ than *cat2* and *cat3* single mutants (Figures 5E and S5). Moreover, *pif4cat2-1* and *pif4cat3* double mutants showed hypocotyl phenotype and H_2_O_2_ levels similar to the *pif4* mutant under high temperature, confirming PIF4 functions upstream to CAT2 and CAT3 for their transcriptional regulation. Conversely, overexpression of *CAT2* in *pif4* mutant (*pif4*/35S:CAT2) showed a significant increase in the hypocotyl elongation and reduction in the H_2_O_2_ levels under high temperature compared with *pif4* mutant (Figure 5C-E). However, *pif4*/35S:CAT3 lines showed limited hypocotyl elongation response compared to *pif4*/35S:CAT2 lines (Figure S5C-E). Together, these findings strongly support that PIF4 functions upstream to CAT2 and CAT3 in the same pathway to maintain H_2_O_2_ levels for controlling hypocotyl elongation under high temperature.

### PIF4 controls CAT2 abundance, whereas H_2_O_2_ modulates PIF4 stability forming a PIF4-CAT-H_2_O_2_ regulatory module

While we established the transcriptional regulation of *CAT*s by PIF4 using molecular and genetic experiments, *CAT*s encode for enzymes. Therefore, we tested whether the PIF4-mediated transcriptional regulation of *CAT*2 correlates with the protein level. Consistent with the gene expression pattern, *pif4* did not show induction of CAT2 protein under high temperature, in contrast to Col-0 seedlings, connecting PIF4 functions to CAT2 enzyme abundance and function via transcriptional regulation for the control of hypocotyl elongation (Figure 6A). Interestingly, H_2_O_2_ is also reported to promote degradation of a target protein by post-translational modification of cysteine residues of the protein through a process called sulfenylation. ^55–58^ PIF4 protein has seven cysteine residues (Cys-37, Cys-168, Cys-208, Cys-209, Cys-247, Cys-261, and Cys-329), out of which Cys-329 is located in the C-terminal bHLH domain of PIF4 protein (Figure S6). Therefore, we tested whether elevated H_2_O_2_ levels can modulate PIF4 protein abundance. 3AT and H_2_O_2_-treated seedlings, with increased H_2_O_2_ levels, showed significantly reduced PIF4 protein abundance compared to mock seedlings under high temperature (Figure 6B). We further verified this genetically using *cat* mutants that accumulate higher H_2_O_2_ levels. Consistent with the pharmacological treatment, *cat2-1* and *cat2-1cat3* mutants showed strongly reduced PIF4 levels and failed to show induction of PIF4 under high temperature compared to the wild-type seedlings (Figure 6C). However, *cat3* mutant maintained higher PIF4 abundance than *cat2-1* and *cat2-1cat3* mutants, although still lower than Col-0 seedlings (Figure 6C), suggesting *CAT2* isoform plays a more prominent role than *CAT3* in maintaining H_2_O_2_ homeostasis under high temperature. These results show that while PIF4 transcriptionally regulates *CAT*s to reduce H_2_O_2_ accumulation, higher H_2_O_2_ levels reduce PIF4 protein abundance post-translationally, suggesting a PIF4-CAT-H_2_O_2_ negative feedback loop that coordinates ROS-homeostasis to control hypocotyl elongation in response to high temperature.

**Figure 6.**
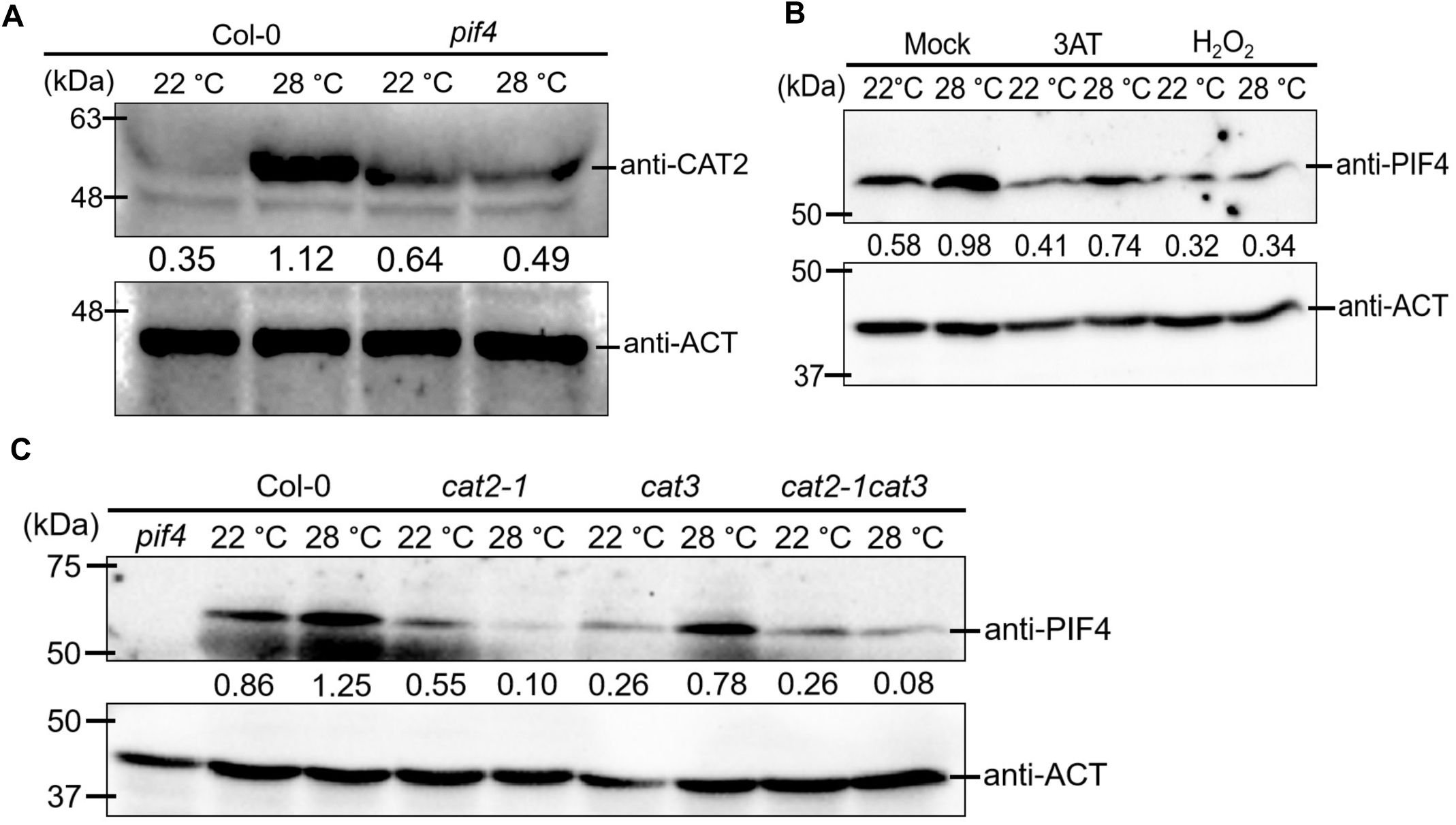
PIF4-mediated regulation of CAT2 protein and H_2_O_2_-mediated control of PIF4 protein abundance. (A) Immunoblot showing CAT2 protein levels in Col-0 and *pif4* mutant under control and high temperature conditions. (B) Immunoblot showing PIF4 protein levels in mock, along with 3AT and H_2_O_2_-treated Col-0 seedlings grown under control and high temperature conditions. (C) Immunoblot showing PIF4 protein levels in Col-0, *cat2-1*, *cat3,* and *cat2-1cat3* seedlings grown under control and high temperature conditions. ACTIN was used as an endogenous control, and numbers mentioned below the CAT2 and PIF4 lanes represent the relative band intensity of CAT2 or PIF4 proteins normalized to ACTIN.

### PIF4-CAT-H_2_O_2_ regulatory module integrates with auxin signaling for the regulation of hypocotyl elongation under high temperature

Next, we investigated the signaling downstream to the PIF4-CAT-H_2_O_2_ regulatory module for controlling hypocotyl elongation under high temperature. Auxin biosynthesis and signaling, along with brassinosteroid signaling, are shown to be critical for high-temperature-mediated hypocotyl elongation. ^13, 15, 18, 20, 59^ Therefore, we examined the expression of auxin biosynthesis and signaling genes that are known to be induced under high temperature conditions in *cat* mutants, along with Col-0 and *pif4* controls. High temperature induced the expression of auxin biosynthetic gene *YUCCA8* in Col-0. However, no induction of *YUCCA8* was observed in *cat2* mutants, similar to *pif4*, in high temperature (Figures S7A). The *cat3* mutant showed attenuated induction of *YUCCA8* in high temperature compared to Col-0. Similarly, high temperature-induced auxin-responsive genes *IAA19* and *SAUR19* also showed no induction in *cat2* and limited induction in *cat3* mutants in comparison with Col-0 (Figure S7B–8C).

Since we observed compromised expression of auxin biosynthesis and signaling genes in *cat* mutants compared to Col-0, attenuated hypocotyl elongation of the *cat* mutants in high temperature might be attributed to reduced auxin levels or signaling. To test this, we supplemented *cat* mutants with exogenous auxin to investigate the effects on hypocotyl elongation. We observed that Picloram (PIC) treatment significantly increased hypocotyl elongation in both Col-0 and *pif4* in control and high temperature conditions compared with mock-treated seedlings (Figure 7A). PIC treatment restored hypocotyl elongation response of *cat2-1*, *cat2-2*, *cat3*, and *cat2-1cat3* mutants, similar to wild-type mock-treated seedlings grown under high temperature (Figure 7A). However, PIC treatment could not completely restore the hypocotyl elongation response of *pif4*, *pif4cat2-1,* and *pif4cat3* seedlings, suggesting an additional level of regulation by PIF4 for hypocotyl elongation under high temperature (Figure 7A). Altogether, these results suggest that the PIF4-CAT-H_2_O_2_ regulatory module for the ROS-homeostasis integrates with auxin biosynthesis and signaling for hypocotyl elongation under high temperature.

**Figure 7.**
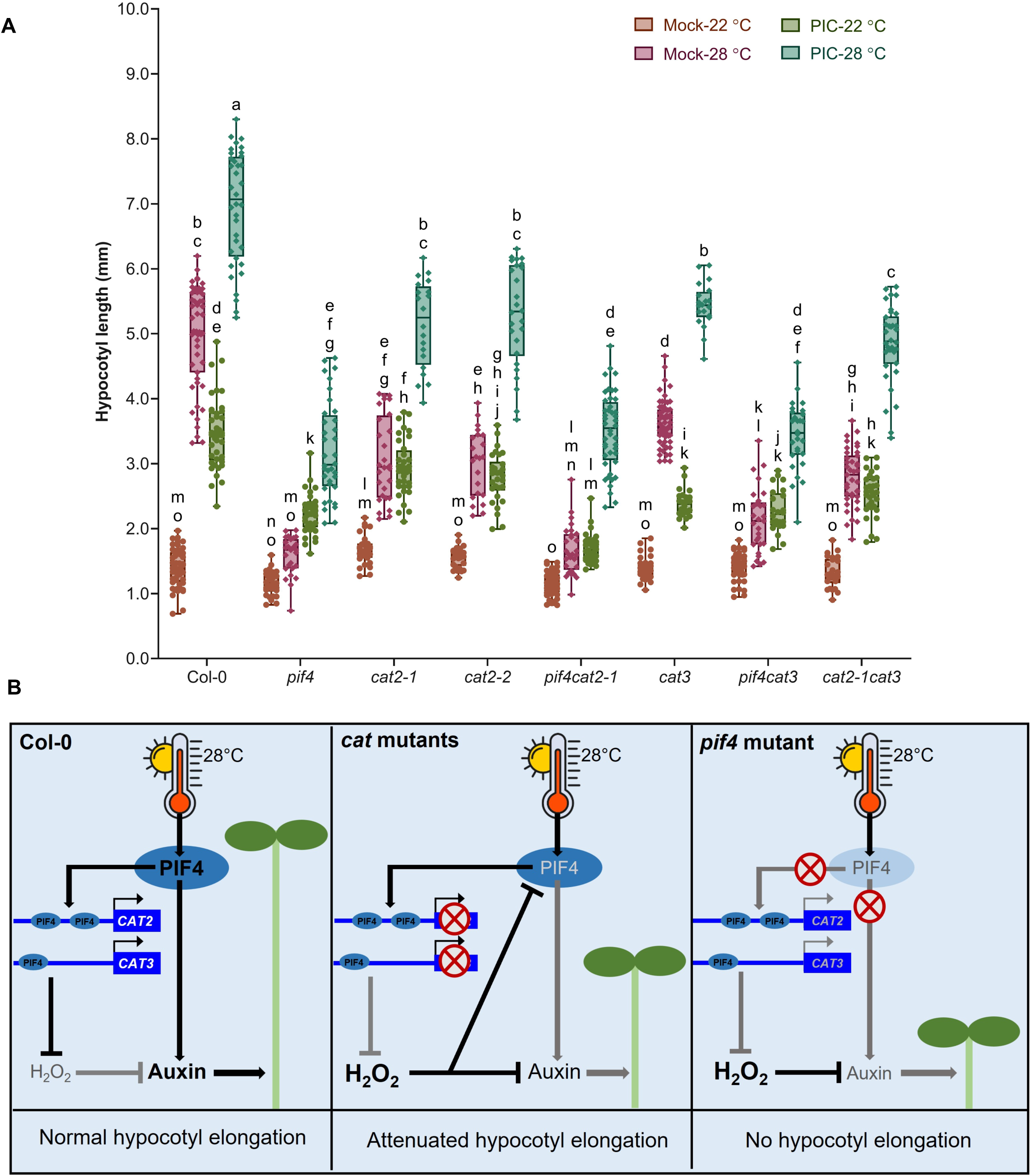
PIF-CAT-H_2_O_2_ regulatory module integrates with auxin signaling for regulating seedling thermomorphogenesis. (A) Box plot showing hypocotyl elongation response of *pif4*, *cat2-1*, *cat2-2*, *pif4cat2-1*, *cat3*, *pif4cat3,* and *cat2-1cat3* mutants along with Col-0 control upon exogenous treatment of Picloram (PIC-1 µM) under control (22 °C) and high temperature (28 °C) conditions. Error bars represent SD from the mean value. Different lowercase letters denote the statistical significance among the samples, as determined by two-way ANOVA followed by Tukey’s post hoc test (*P*<0.01). (B) Schematic diagram showing PIF4-CAT-H_2_O_2_ regulatory module-mediated regulation of thermomorphogenic hypocotyl elongation in Arabidopsis. High temperature induces PIF4, which in turn transcriptionally activates *CAT2* and *CAT3* genes to maintain H_2_O_2_ levels in Col-0. In parallel, PIF4 directly regulates auxin biosynthesis and signaling to control hypocotyl elongation. In *cat* mutants, elevated H_2_O_2_ levels attenuate auxin signaling to reduce hypocotyl length under high temperatures. Elevated H_2_O_2_ level further reduces the PIF4 protein abundance, leading to impaired PIF4-auxin module to attenuate hypocotyl elongation. In the *pif4* mutant, loss of PIF4 function compromises both CAT-mediated H_2_O_2_ homeostasis and PIF4-dependent regulation of auxin biosynthesis and signaling, which results in failure of hypocotyl elongation in high temperature. Bold font denotes higher protein, H_2_O_2,_ or auxin levels, whereas solid black and grey lines represent strong and weaker regulatory interactions, respectively. Together, the PIF4-CAT-H_2_O_2_ regulatory module integrates H_2_O_2_ homeostasis with auxin signaling to coordinate hypocotyl elongation response in high temperature conditions.

## Discussion

Plants exhibit remarkable developmental plasticity under high temperatures throughout their life cycle.^1–3, 60, 61^ Hypocotyl elongation is one of the important adaptive responses at the seedling stage to help seedlings grow in high temperatures. Genetic regulation of hypocotyl elongation under high temperature involves thermoreceptors, transcription factors, such as PIF4, and phytohormones, such as auxin. While ROS functions as one of the key signaling molecules regulating plant developmental and physiological responses under environmental changes and various stresses, including heat stress, the involvement of ROS-homeostasis in regulating thermomorphogenesis remained elusive. In this study, we demonstrate that the homeostasis of H_2_O_2_, the most stable ROS, is essential for seedling thermomorphogenesis. Our study reveals an interplay between temperature signaling and ROS signaling, where PIF4 plays a key role in maintaining H_2_O_2_ homeostasis through transcriptional activation of *CAT2* and *CAT3* genes (Figure 7B). Moreover, elevated H_2_O_2_ levels reduce the PIF4 protein abundance to fine-tune the seedling thermomorphogenesis. The deciphered PIF4-CAT-H_2_O_2_ regulatory module eventually integrates to auxin biosynthesis and signaling for regulating the hypocotyl elongation under high temperature.

While temperature-induced changes in ROS levels were expected, systematic meta-analysis of the available transcriptomic datasets for the Arabidopsis seedlings exposed to high temperature for different time points ^15, 22, 52^ combined with our hypocotyl-specific transcriptome analysis for high temperature suggested a strong effect of high temperature on ROS signaling and homeostasis (Figure 1). A recent report, however, showed that 2-hour treatment of high temperature (27 °C) and heat stress (37 °C) showed no significant changes in O_2_^•−^ and H_2_O_2_, as detected by NBT and DAB staining, respectively, in the cotyledons and hypocotyl of Arabidopsis seedlings. ^22^ In contrast, 4-hour shade treatment increased the O_2_^•−^ and H_2_O_2_ accumulation, leading to altered redox balance in the hypocotyl cells to promote elongation.^44^ Our time course gene expression analysis, combined with DAB staining for H_2_O_2_ accumulation in hypocotyls, unveiled 12 hours as the crucial time point for genetic changes associated with ROS levels in response to high temperature (28 °C). While high ambient temperature induces the expression of ROS-related genes, ROS homeostasis, and not the absolute ROS levels, is shown to be critical in a developmental context. ^33, 35, 62, 63^ For example, ROS homeostasis is crucial for the transition of tapetal cells from cell division to cell differentiation in anthers. ^33^ Similarly, disruption of ROS homeostasis impairs root development by affecting lateral root emergence, root primordia, and root hair growth. ^35, 64–67^ Our results support that maintaining proper ROS levels, instead of a strong induction, is required for optimal hypocotyl elongation response under high temperature. Increasing the H_2_O_2_ levels beyond a threshold would rather inhibit temperature-mediated hypocotyl elongation (Figure 2). The function of the enzymes involved in ROS production and scavenging is key to maintaining the optimal ROS levels. ^39, 54^ Catalases, the key enzymes for H_2_O_2_ scavenging that maintain peroxisomal H_2_O_2_ levels in response to various environmental cues, would be expected to be crucial for maintaining optimal H_2_O_2_ levels for thermomorphogenesis. ^35, 54^ This was supported by the induction of *CAT2* and *CAT3* at 12 hours post high temperature treatment, the critical time point for genetic changes related to ROS signaling in response to high temperature, coinciding with the induction of several ROS-metabolism genes. Consistent with that, *cat* mutants as well as *CAT* overexpression lines showed altered hypocotyl elongation than wild-type seedlings under high temperature. Interestingly, both *cat* mutants, with significantly higher H_2_O_2_ levels than wild-type seedlings, and *CAT* overexpression lines, with remarkably lower H_2_O_2_ levels, showed attenuated hypocotyl elongation under high temperature, strengthening the requirement of CAT-mediated H_2_O_2_ homeostasis for seedling thermomorphogenesis (Figure 3).

Transcriptional regulation, involving bHLH transcription factor PIF4, is an integral component of Arabidopsis thermomorphogenesis. Temperature-induced PIF4 binds to the promoters of auxin biosynthesis and responsive genes, as well as promotes brassinosteroid (BR) biosynthesis to regulate hypocotyl elongation. ^16–19^ Moreover, transcriptional regulation of the antioxidant enzymes is suggested to be a primary mechanism to maintain the H_2_O_2_ level. For example, a Myb-like transcription factor KUODA1 (KUA1) binds to the promoters of genes encoding Peroxidases (Prxs), reducing H_2_O_2_ levels to promote cell expansion in Arabidopsis leaves. ^62–63^ SPEECHLESS (SPCH), bHLH transcription factor, binds to the promoters of *CAT2* and *APX1*, encoding two major antioxidant enzymes, to regulate H_2_O_2_ levels for controlling stomatal development. ^30^ Maintaining optimal H_2_O_2_ levels for thermomorphogenesis would require integration of temperature signaling with ROS signaling for the regulation of downstream phytohormonal response and cellular attributes. Indeed, PIF4, a key transcriptional regulator of temperature signaling, binds to the promoters of *CAT2* and *CAT3* genes, encoding major enzymes of H_2_O_2_ scavenging, under high temperature to maintain optimal H_2_O_2_ levels for promoting hypocotyl elongation (Figure 4). Failure of *pif4* mutant to elongate hypocotyl under high temperature could, at least in part, be attributed to inefficient H_2_O_2_ scavenging and strong H_2_O_2_ accumulation due to reduced activation of CATs (Figure 5). Partial rescue of the *pif4* hypocotyl phenotype under high temperature by promoting H_2_O_2_ scavenging, either by exogenous DMTU treatment or by overexpressing *CAT*, further supports PIF4-mediated integration of temperature and ROS signaling to maintain optimal H_2_O_2_ levels. Thus, PIF4 functions upstream to CATs to form an important module to control seedling thermomorphogenesis by regulating H_2_O_2_ homeostasis. This PIF4-CAT regulatory module appears to function in parallel to the well-established PIF4-mediated direct regulation of auxin biosynthesis for hypocotyl elongation under high temperature, as *cat* mutants have a weaker hypocotyl elongation phenotype than *pif4* mutants, and CAT overexpression only partially rescues the *pif4* hypocotyl phenotype under high temperature.

Our findings show reduced PIF4 protein abundance due to increased H_2_O_2_ levels under high temperature, thereby providing an additional regulatory switch to control seedling thermomorphogenesis via integration of ROS and temperature signaling. Genetic mutation of *CAT2* leads to a strong reduction in PIF4 abundance due to higher H_2_O_2_ accumulation, whereas *CAT3* mutation results in higher PIF4 abundance than *cat2* but lesser than wild-type seedlings, suggesting dominance of CAT2 over CAT3 to maintain H_2_O_2_ homeostasis to preserve PIF4 stability and ensure proper thermomorphogenic response (Figure 6). Protein homeostasis usually involves multiple interconnected levels of regulation, such as transcriptional, translational, and post-translational regulations. These multilayered controls ensure proper cellular functioning and developmental responses to adapt to changing environmental conditions. ^18, 68–70^ We speculate that H_2_O_2_-mediated regulation of PIF4 protein abundance is likely due to sulfenylation, as H_2_O_2_ is documented to modify cysteine amino acids of a target protein by converting thiol (-SH) to sulphenic acid (-SOH) group that leads to degradation of the target protein. ^55–58, 71–72^ The presence of seven cysteine residues (Cys-37, Cys-168, Cys-208, Cys-247, Cys-261, Cys-329, and Cys-209) in the PIF4 protein that are amenable to sulfenylation, one of which (Cys-329) is located in the C-terminal bHLH domain of the protein, further supports the hypothesis. This would allow H_2_O_2_ to keep a check on PIF4 protein levels, while PIF4 is keeping a check on H_2_O_2_ via transcriptional activation of *CAT*s. Together, we propose a PIF4-CAT-H_2_O_2_ negative feedback loop for maintaining H_2_O_2_ homeostasis to regulate Arabidopsis seedling thermomorphogenesis.

Our study established a novel PIF4-CAT-H_2_O_2_ regulatory module, with transcriptional as well as translational control, that modulates hypocotyl elongation under high temperature. However, the signaling downstream to the PIF4-CAT-H_2_O_2_ module for regulating hypocotyl length was intriguing. Auxin biosynthesis and signaling are integral components of temperature-mediated hypocotyl elongation downstream of PIF4. Moreover, ROS have been reported to influence auxin homeostasis by affecting both its biosynthesis and signaling. ^44, 73–74^ *cat2* mutant, with less auxin levels, showing hyponastic leaves under high light conditions that get rescued with exogenous auxin supplementation, indicated that the PIF4-CAT-H_2_O_2_ regulatory module may also integrate to auxin signaling for hypocotyl elongation. ^75^ Expression of auxin biosynthesis and signaling genes in *cat* mutants compared with wild-type seedlings, as well as the complete rescue of the attenuated hypocotyl elongation phenotype of *cat* mutants by exogenous auxin supplementation, confirmed the integration of the PIF4-CAT-H_2_O_2_ module to auxin signaling for the regulation of seedling thermomorphogenesis. Mechanistically, H_2_O_2_-induced depletion of auxin level could be attributed to H_2_O_2_-mediated sulfenylation that suppresses the activity of TRYPTOPHAN SYNTHETASE β SUBUNIT 1 (TSB1), leading to depletion in the level of tryptophan, a precursor molecule required for auxin biosynthesis. ^55^ Thus, in addition to the established direct transcriptional regulation of auxin biosynthesis and signaling by PIF4, our findings support an additional pathway in which PIF4 controls auxin biosynthesis and signaling through CAT-mediated modulation of H_2_O_2_ levels to fine-tune the hypocotyl elongation under high temperature. Complete rescue of *pif4* hypocotyl phenotype by overexpressing *SMALL AUXIN UP RNA 19* (*SAUR19*) under high temperature^17^ suggests that auxin acts as a downstream player in PIF4-mediated control of thermomorphogenesis via both direct regulation of auxin biosynthetic genes and H_2_O_2_-homeostasis-mediated regulation of auxin biosynthesis and signaling.

In summary, our study shows that the high temperature promotes PIF4-mediated transcriptional activation of *CAT2* and *CAT3* genes, thereby maintaining optimal H_2_O_2_ homeostasis for proper hypocotyl elongation. Genetic and pharmacological experiments showed that PIF4 and CATs work together in the same pathway, and PIF4 functions upstream of CATs. Moreover, elevated H_2_O_2_ levels negatively affect PIF4 protein abundance, providing an additional level of regulation for maintaining hypocotyl growth in high temperatures. The identified PIF4-CAT-H_2_O_2_ negative feedback loop integrates to auxin biosynthesis and signaling to drive hypocotyl elongation. Collectively, our findings show the interplay of temperature signaling and ROS signaling that converge on auxin biosynthesis and signaling for the regulation of Arabidopsis seedling thermomorphogenesis (Figure 7B).

## Materials and Methods

### Plant material, growth conditions

The Columbia-0 (Col-0) ecotype of *Arabidopsis thaliana* was used in the present study. Various T-DNA insertion lines (mutants) of *cat2-1* (CS67045), *cat2-2* (SALK_076998), *cat3* (SALK_209162C), *pif4-2* (CS66043), and *proPIF4*:GUS (CS69169) used in the present study were procured from the Arabidopsis Biological Resource Center (TAIR). Homozygous double cross mutants of *cat2-1cat3*, *cat2-1pif4*, and *cat3pif4* were generated by crossing. Additional lines such as *35S:CAT2*, *35S:CAT3*, *proCAT2:GUS*, *proCAT3:GUS*, *pif4*/*proCAT2:GUS*, *pif4*/*proCAT3:GUS*, *pif4*/*35S:CAT2,* and *pif4*/*35S:CAT3* lines were generated using the floral dip transformation method. ^76^

Arabidopsis seeds were surface sterilized and stratified in the dark at 4 °C for 48 h. After stratification, seeds were sown on half-strength Murashige and Skoog (MS) medium plates, pH 5.7, containing 0.8% Agar (w/v). Seeds were germinated and seedlings were grown for 4 days under 100 µmol m^−2^ s^−1^ light intensity with a 16 h day/8 h night photoperiod in a growth room maintained at 22 °C. For high temperature phenotyping, plates were transferred to 28 °C with the same light conditions, and seedlings were grown for the next 72h. For pharmacological experiments, seedlings were transferred to half-strength MS medium supplemented with ROS scavenger N, N’-Dimethylthiourea (DMTU, 0.25 to 5.0 mM), catalase inhibitor 3-Amino-1,2,4-triazole (3AT, 0.05 to 5.0 µM), and synthetic auxin analogue Picloram (PIC, 1.0 µM) on the 4^th^ day and were grown for the next 72h either at 22 °C or 28 °C. Change in hypocotyl length was measured on the 7^th^ day after sowing and compared with the seedlings grown at the control (22 °C) condition.

### Hypocotyl length quantification

To quantify the hypocotyl length, 40-50 seedlings were used for each genotype in different conditions and treatments. Seedlings were photographed using a digital camera (Sony), and the digital images were used to measure hypocotyl lengths of seedlings using ImageJ version 1.54g (https://imagej.net/ij/download.html). Representative photographs of the seedlings of each genotype for hypocotyl elongation phenotype were taken using a stereo microscope (SZM167D, LMI, UK).

### Meta-analysis of publicly available transcriptome data

We used RNA-seq data from three independent studies representing short- to long-term high temperature treatments (1hr, 2hrs, 3hrs, 6hrs, 24hrs, and 48hrs) (Burko et al., 2022; Sang et al., 2023; Dard et al., 2023). To ensure comparability across datasets, raw gene expression counts from each study were normalized individually. Normalization was performed using the Trimmed Mean of M-values (TMM) method in the edgeR package. ^77^ Differentially expressed genes (DEGs) were identified using the exactTest() function, comparing high temperature vs. control groups. Significance was determined using Benjamini-Hochberg correction for false discovery rate (FDR). DEGs were defined as genes with FDR < 0.05 and log_2_FC values ≥ 0.585 or ≤ -0.585. Gene Ontology (GO) enrichment analysis was performed for upregulated genes at different points using the Plant Transcriptional Regulatory Map (PlantRegMap). ^78^ GO terms with *P*-value< 0.005 and gene count more than 5 were selected. Bubble plots for enriched GO terms were prepared using the ggplot2 package in R.

### RNA extraction and gene expression studies

Seedlings without roots (cotyledons and hypocotyls) were considered for gene expression studies. Seedlings were grown for 6 days at 22 °C and then were transferred to high temperature (28 °C) for different time-points (from 1hr to 24hrs). At each time point, the aerial part of the seedling was cut using micro scissors, immediately frozen in liquid nitrogen, and stored in –80°C. Total RNA was extracted using QIAzol lysis reagent (Qiagen), and 2.0 µg total RNA was considered for cDNA synthesis using Verso cDNA Synthesis Kit (Thermo Fisher Scientific). For the qPCR experiment, cDNA was diluted with nuclease-free water and used as a template. Gene-specific primers and PowerUp™ SYBR™ Green master mix (Applied Biosystems) were used for performing qPCR in a Bio-Rad CFX9 connect thermocycler system (Bio-Rad). Transcript abundance of each gene was represented relative to the level of 18S (*AT3G41768*) or GAPDH (*AT1G13440*) using the 2^−ΔCT^ method. ^79–80^ For each qPCR experiment, two biological replicates, each having three technical replicates (n=6), were considered. Primer pairs used to check the expression of various genes in this study are presented in **Table S1**.

### cDNA library preparation and RNA-seq analysis

6-days-old Arabidopsis seedlings grown at 22 °C were shifted to 28 °C for 1 hour treatment at ZT1. Hypocotyls of Arabidopsis seedlings were dissected and snap-frozen in liquid nitrogen. The samples were collected in three biological replicates. Libraries were prepared in three biological replicates using YourSeq Full Transcript RNA-seq Library Kit (Amaryllis Nucleics, Oakland, CA, USA) following the manufacturer’s protocol. These RNA-seq libraries were sequenced on the Illumina HiSeq 4000 platform to generate 150-bp paired-end reads. The quality-filtered reads were mapped to the representative cDNA sequence of *Arabidopsis thaliana* (TAIR10) obtained from Phytozome (https://phytozome-next.jgi.doe.gov/) using the default parameters of Bowtie2. ^81^ Differential gene expression analysis was performed by doing pairwise comparisons using the edgeR package. ^77^ The analysis utilized the glm approach with a quasi-likelihood F-test. Significantly differentially expressed genes in each pair-wise comparison were identified using an FDR cut-off < 0.05 and logFC values ≥ 0.585 or ≤ –0.585. Gene ontology analysis was performed utilizing the GO enrichment tool of the Plant Transcriptional Regulatory Map. ^78^ GO terms with q-value<0.05 and gene count more than 10 were considered to be significantly enriched. Bubble plots for the enriched GO terms were prepared using the ggplot2 package in R.

### DAB staining and H_2_O_2_ quantification

Arabidopsis seedlings were grown at 22 °C for 6 days and shifted to 28 °C for 12 h. After that, control and treated seedlings were incubated in freshly prepared DAB staining solution [3, 3’- Diaminobenzidine (1 mg/ml), Tween 20 (0.05% v/v), Na_2_HPO_4_ (10 mM)] at room temperature in the dark for 6-8 hrs. After that DAB staining solution was discarded, and a bleaching solution (ethanol: acetic acid: glycerol in a ratio of 3:1:1 v/v/v) was used to remove pigments from the stained seedlings. DAB-stained seedling of each genotype was photographed using a stereo microscope (SZM167D, LMI, UK). To quantify the H_2_O_2_ levels, stained seedlings were ground in liquid nitrogen and resuspended in 0.2 M HClO4 and centrifuged at 10,000 x g for 10 mins at 4 °C. The supernatant was used to measure the absorbance at 450 nm, and H_2_O_2_ was quantified using a standard curve, plotted with known concentrations of H_2_O_2_ in 0.2 M HClO_4_-DAB.

### Cloning, construct preparation, and genetic transformation

To generate different transgenic lines used in the present study, coding sequences (CDS) and promoters of the selected genes were amplified with their respective primer pairs (**Table S1**) using high-fidelity Phusion polymerase (Thermo Fisher Scientific). Various entry clones of CDS or promoters were made using pENTR/D-TOPO cloning Kit (Invitrogen, Thermo Fisher Scientific). After confirming the cloning with Sanger sequencing, CDS or promoters were cloned into destination vectors using Gateway LR Clonase II enzyme mix (Invitrogen, Thermo Fisher Scientific). These constructs were then mobilized into the *Agrobacterium tumefaciens* strain, GV3101, using the freeze-thaw method. ^82^ After confirming the presence of constructs in the *Agrobacterium* by PCR, genetic transformation of wild-type (Col-0) plants was done using the floral-dip transformation method. ^76^

### Luciferase trans-activation and Dual luciferase assay

To check the interaction of transcription factor with the promoters of the target genes, luciferase trans-activation assay and dual luciferase assays were performed. Upstream promoter region of *CAT2* (3,022-bp) and *CAT3* (1,882-bp) genes were cloned into the pGWB435 gateway binary vector to generate *proCAT2:LUC* and *proCAT3:LUC* constructs, respectively. Similarly, CDS of PIF4 (1290-bp) was cloned into the pGWB402 gateway binary vector to generate *35S:PIF4* construct. Agrobacterium strain, GV3101, carrying these constructs, were co-infiltrated in *N. benthamiana* leaves in four different combinations: (1) infiltration buffer, (2) mixture of empty vectors pGWB435 + pGWB402, (3) promoter:LUC + empty vector pGWB402, (4) promoter:LUC + 35S:PIF4, at 4 different spots. After 48h of infiltration, the infiltrated leaves were sprayed with D-Luciferin, potassium salt (GOLDBIO), and luminescence was detected using the ChemiDoc MP imaging system (Bio-Rad). At least 10 different infiltrated leaves were considered to conclude the interaction results. For the dual luciferase assay, dual-pro*CAT2*:LUC and dual-pro*CAT3*:LUC constructs were made using another gateway binary vector p635nRRF, having *35S:REN* (Renilla luciferase) as an internal control. To quantify the relative luciferase activity (LUC/REN ratio), constructs dual-pro*CAT2*:LUC, dual-pro*CAT3*:LUC were infiltrated in *N. benthamiana* leaves with and without *35S:PIF4*. Leaf discs (1 cm diameter) from the site of infiltration were cut, and leaf extracts were used to measure Firefly and Renilla luciferase activities using the Dual-luciferase assay system (Promega, Madison, WI, USA) according to the manufacturer’s protocol. Luminescence was quantified using a luminometer POLARstar Omega (BMG Labtech).

### GUS staining and quantification

Control seedlings grown at 22 °C and high temperature-treated seedlings (exposed to 28 °C for 12 h) were incubated in GUS staining buffer, composed of phosphate buffer (pH-7.0), 0.1% Triton X-100 (v/v), 0.5 mM K_4_Fe(CN)_6_.3H_2_O, 0.5-mM K_3_Fe(CN)_6_ with 958 µM X-gluc substrate, for 6-8 h at 37 °C. Seedlings were then washed with 80% ethanol to remove all pigments, stored in ethanol:acetic acid solution (3:1, v/v), and photographed using a stereo microscope (SZM167D, LMI, UK).

### Chromatin immunoprecipitation (ChIP) assay

ChIP assay was performed following the protocol outlined by ^83, 84^ with minor modifications. In brief, seven-day-old seedlings (both control and temperature-treated) were incubated in cross-linking buffer for 10 min, followed by the addition of 2M glycine to stop the cross-linking reaction. Samples were then washed three times with sterile water, dried with sterile blotting paper, and ground into fine powder. The powder was dissolved in nuclei isolation buffer and centrifuged at 11000 x g for 20 min. The resulting nuclei pellet was resuspended in nuclei lysis buffer, followed by shearing of chromatin into small fragments (500 bp) using a sonicator. The sonicated sample was then centrifuged at 13800 x g for 10 min at 4 °C to separate the debris. Immunoprecipitation was performed using polyclonal rabbit anti-PIF4 antibody (AS16 3157; Agrisera), followed by reverse cross-linking, protein digestion, and DNA elution as mentioned in the protocol. ChIP-qPCR was performed using ∼50 ng DNA template with G-box flanking primers as mentioned in **Table S1**.

### Protein extraction and western blot

∼100 mg each of 7-days-old control seedlings grown at 22 °C and high temperature-treated seedlings (exposed to 28 °C for 12 h) were harvested and total protein was extracted using YODA protein extraction buffer, containing 50 mM Tris pH-7.5, 150 mM NaCl, 1 mM EDTA, 10% glycerol (v/v), 5.0 mM DTT, 1% protease inhibitor cocktail (v/v), 10 mM MG132, 2.0 mM phenylmethylsulfonyl fluoride (PMSF), and 0.1% Triton X-100 (v/v). Protein samples were resolved by SDS-PAGE and transferred to a PVDF membrane (0.45 µm pore size) using the mini blot module (Invitrogen). The membranes were probed with primary antibodies specific to target proteins, followed by a secondary antibody incubation. Protein bands were visualized through a chemiluminescence reaction using Clarity Western ECL substrate (Bio-Rad), with detection performed on a ChemiDoc XRS+ (Bio-Rad) under high-sensitivity chemiluminescence settings. We used polyclonal rabbit anti-CAT2 (AS21 4531, Agrisera), anti-PIF4 (AS16 3157; Agrisera), and anti-ACT (AS13 2640, Agrisera) primary antibodies and goat anti-rabbit IgG (H&L), HRP conjugated (AS09 602, Agrisera) secondary antibody for western blot.

## Accession numbers

PIF4 (*AT2G43010*), PIF5 (*AT3G59060*), PIF7 (*AT5G61270*), CAT2 (*AT4G35090*), CAT3 (*AT1G20620*), CAT1 (*AT1G20630*), SOD1 (*AT1G08830*), SOD2 (*AT2G28190*), MnSOD1 (*AT3G10920*), CSD3 (*AT5G18100*), FSD3 (*AT5G23310*), CSOD4 (*AT1G12520*), FSD1 (*AT4G25100*), FSD2 (*AT5G51100*), MSD2 (*AT3G56350*), RBOH-A (*AT5G07390*), RBOH-B (*AT1G09090*), RBOH-C (*AT5G51060*), RBOH-D (*AT5G47910*), RBOH-E (*AT1G19230*), RBOH-H (*AT5G60010*), RBOH-I (*AT4G11230*), RBOH-J (*AT3G45810*), APX1 (*AT1G07890*), BZR1 (*AT1G75080*), IAA29 (*AT4G32280*), UBC30 (*AT5G56150*), YUCCA8 (*AT4G28720*), IAA19 (*AT3G15540*), SAUR19 (*AT5G18010*), GAPDH (*AT1G13440*), 18S (*AT3G41768*).

## Statistical Analysis

Details of the statistical analysis for each experiment are mentioned in the respective figure legends. RNA-seq experiment was performed with three biological replicates. For qPCR analysis, two independent biological replicates, each having three technical replicates (n=6), were analyzed. All experiments were repeated 2-3 times to ensure reproducibility. Statistical significance was assessed using Student’s t-test for comparisons between two groups and two-way ANOVA for comparisons involving multiple groups. Both tests were performed using GraphPad PRISM (10.3.0).

## Acknowledgments

This work was supported by the Core Research Grant (CRG/2020/002179) from Anusandhan National Research Foundation (ANRF), Department of Science and Technology, India, as well as core funding from the BRIC-National Institute of Plant Genome Research. KS acknowledges SERB-NPDF fellowships (PDF/2018/002387). MKP and AD acknowledge their CSIR-JRF and UGC-JRF, respectively. We thank NIPGR Central Instrumentation Facility (CIF) for their support. We thank Professor Christian Fankhauser, University of Lausanne, for providing us with Arabidopsis *pif457* triple mutant seeds used in the study.

## Author contributions

R.K.R.-Conceptualization, investigation, data curation, visualization and validation, writing-original draft, K.S.-Conceptualization, writing-review and editing, M.K.P-investigation, data curation and visualization, formal analysis, A.D.- investigation, data curation and visualization, formal analysis, A.R.-Conceptualization, project administration and supervision, funding acquisition, writing-review and editing. All authors have read and approved the final version of the manuscript.

## Supplementary Materials

### Supplemental Datasets

**Supplemental Dataset 1.** List of significant differentially expressed genes in Arabidopsis hypocotyl on 1-hour high ambient temperature treatment

**Supplemental Dataset 2.** Enriched gene ontology categories for differentially expressed genes in Arabidopsis hypocotyl on 1-hour high ambient temperature treatment

### Supplemental Figures

**Figure S1.**
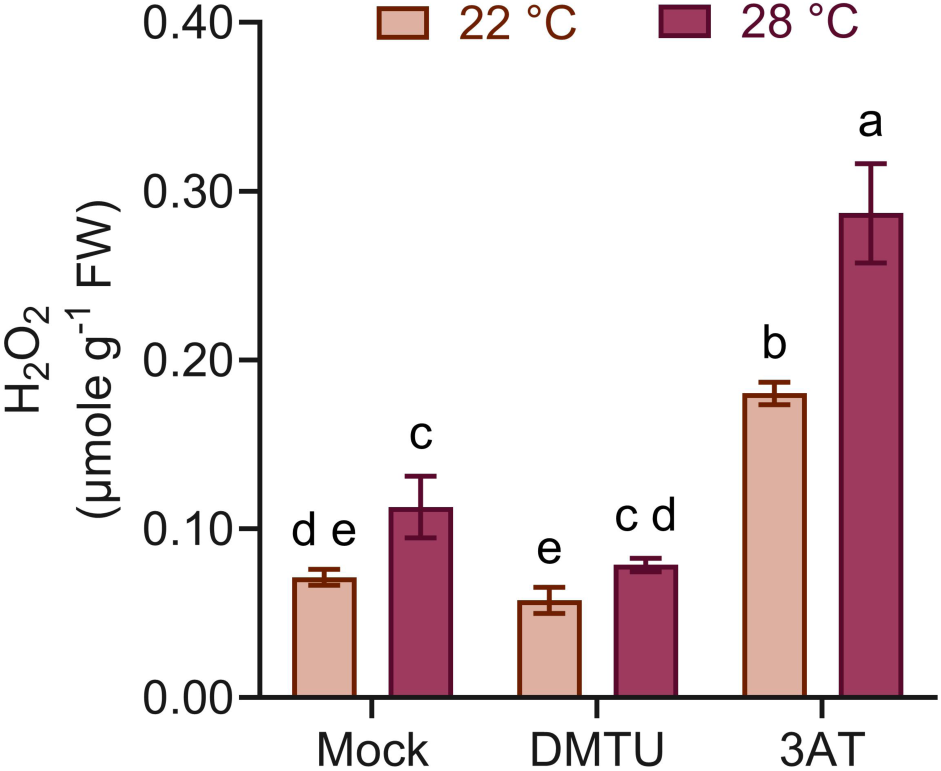
H_2_O_2_ quantification in the seedlings grown on mock, DMTU (2.5 mM)- and 3AT (2.5 μM)- supplemented media. Error bars represents SD. Different lowercase letters above the bars indicate statistically significant differences between the samples, as determined by two-way ANOVA followed by Tukey’s post hoc test (*P*<0.01).

**Figure S2.**
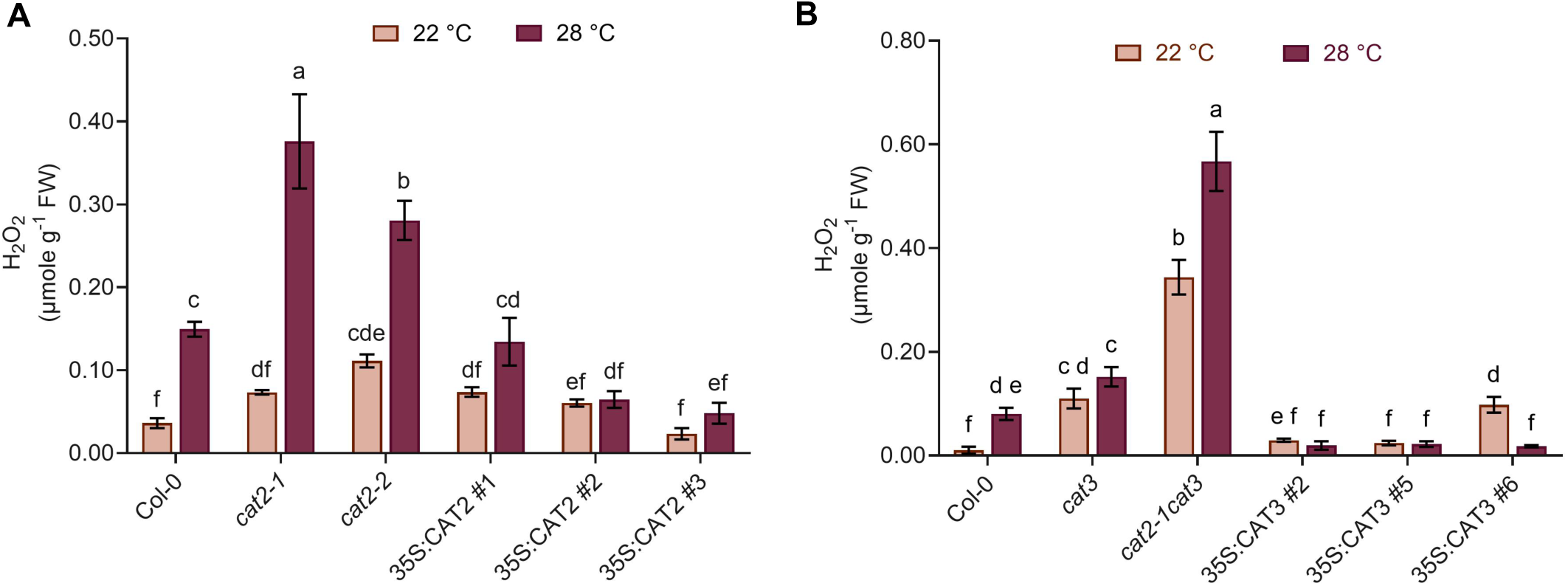
CAT2 and CAT3 control H_2_O_2_ levels under high temperature. (A) Quantification of H_2_O_2_ levels in *cat2* mutants and *35S:CAT2* over expression lines grown under control and high temperature conditions (B) Quantification of H_2_O_2_ levels in *cat3* and *cat2-1cat3* mutants along with *35S:CAT3* over expression lines grown under control and high temperature conditions. Error bar represents SD. Different lowercase letters above the error bars indicates statistically significant differences between the samples, as determined by two-way ANOVA followed by Tukey’s post hoc test (*P*<0.01).

**Figure S3.**
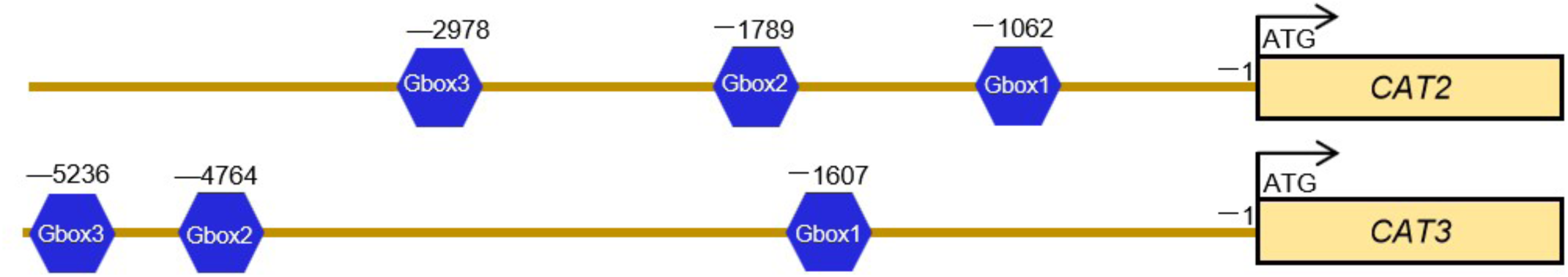
Schematic representation of the positions of PIF4 binding sites in *CAT2* and *CAT3* promoters. G-box (5’-CACGTG-3’) located in the in the upstream promoter of *CAT2* and *CAT3* genes are shown, where negative values represent upstream distances from the start codon ATG of the gene.

**Figure S4.**
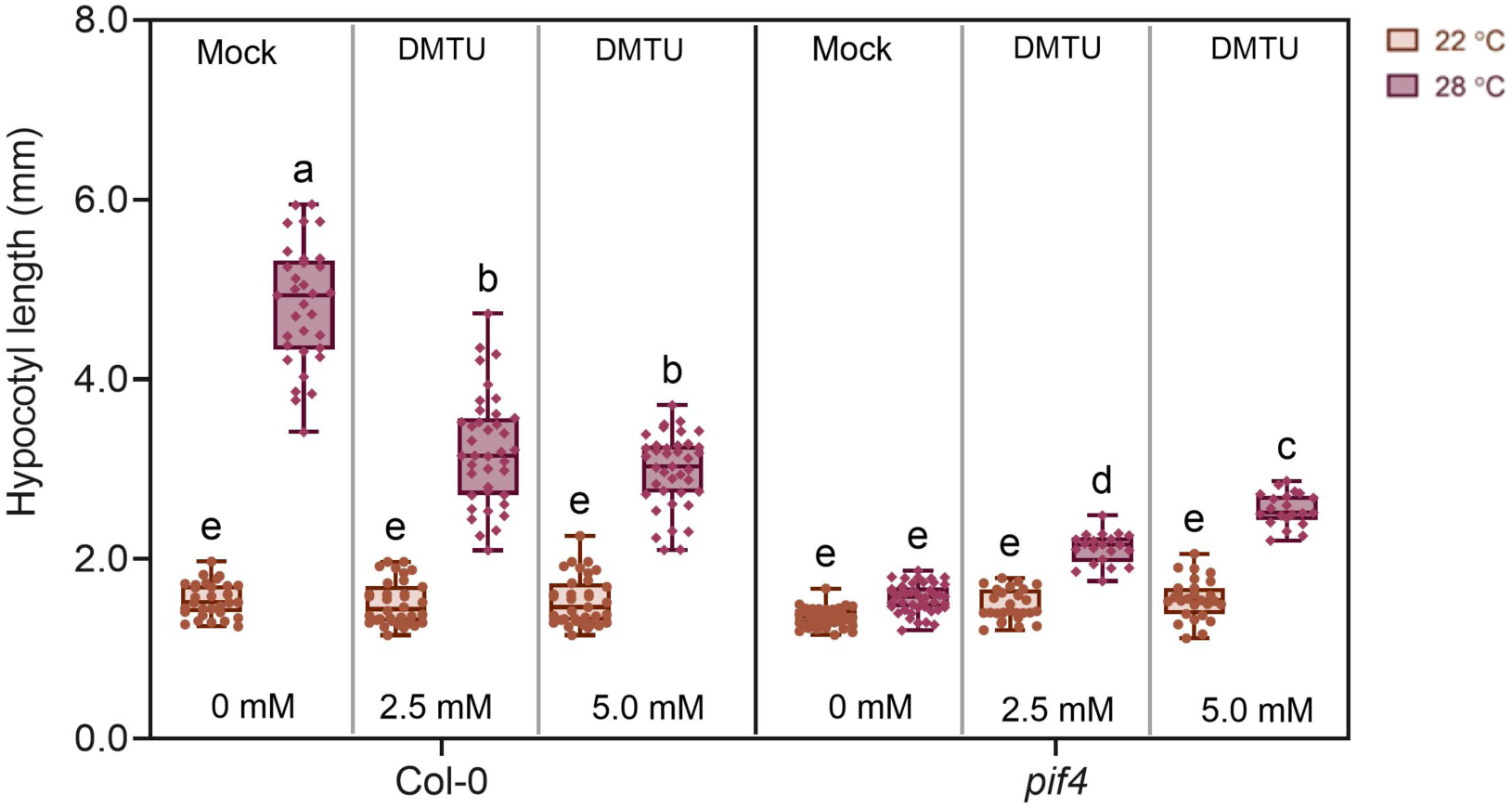
H_2_O_2_ scavenging partially rescues the hypocotyl elongation response of the *pif4* mutant. Hypocotyl elongation response of Col-0 and *pif4* mutant seedlings grown under control and high temperature conditions with exogenous DMTU treatments (2.5 and 5.0 mM). Central horizontal lines within the each box represents the mean. Small circles and diamonds on the box represent number of seedlings quantified. Different lowercase letters indicate statistically significant differences among the samples, as determined by two-way ANOVA followed by Tukey’s post hoc test (*P*<0.01).

**Figure S5.**
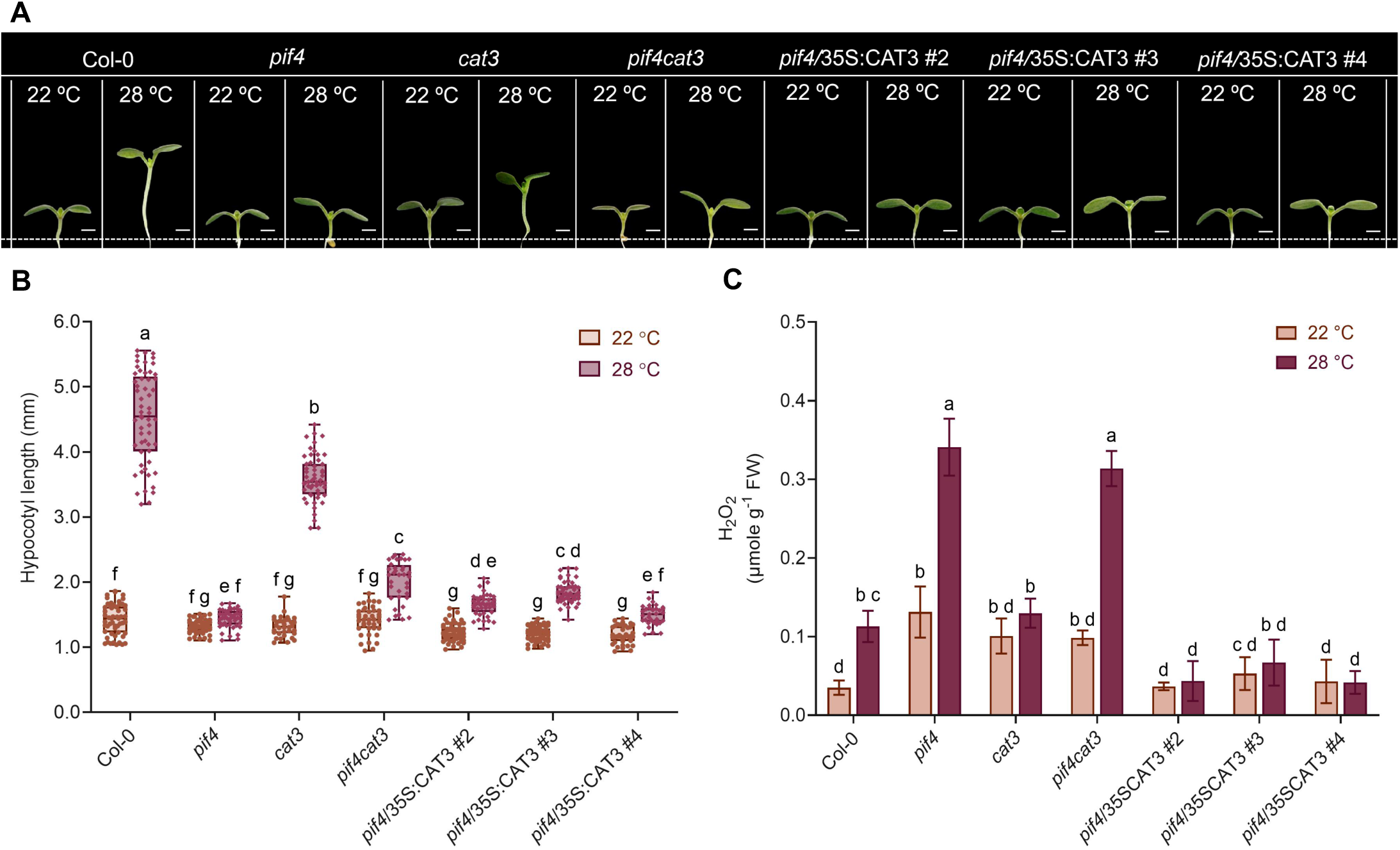
Genetic interaction of *PIF4* and *CAT3* to control seedling thermomorphogenesis. (A) Representative seedling images showing hypocotyl elongation of Col-0, *pif4*, *cat3*, and *pif4cat3* double mutant, along with three independent *pif4*/35S:CAT3 lines grown under control and high temperature conditions. scale bar - 1 mm. (B) Quantification of hypocotyl length of the genotypes shown in (A). (C) Quantification of H_2_O_2_ levels in the genotypes shown in (A) in control and high temperature conditions. Error bars in (B) and (C) represent the SD from the mean values. Different lowercase letters indicate statistically significant differences among the samples, as determined by two-way ANOVA followed by Tukey’s post hoc test (*P*<0.01).

**Figure S6.**
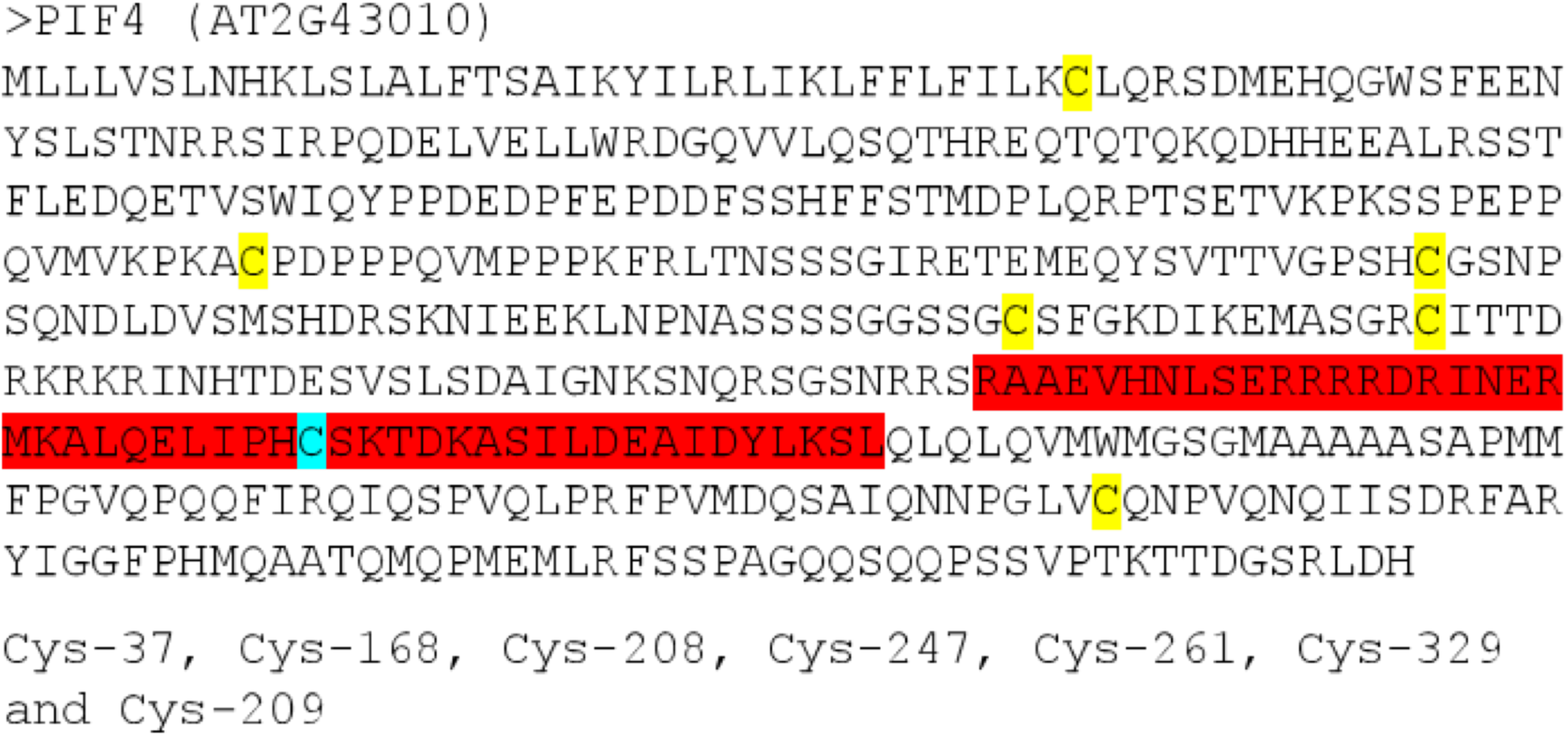
Peptide sequence of PIF4 where with Cysteine (C) residues highlighted. Sequence highlighted in red color represents bHLH-domain.

**Figure S7.**
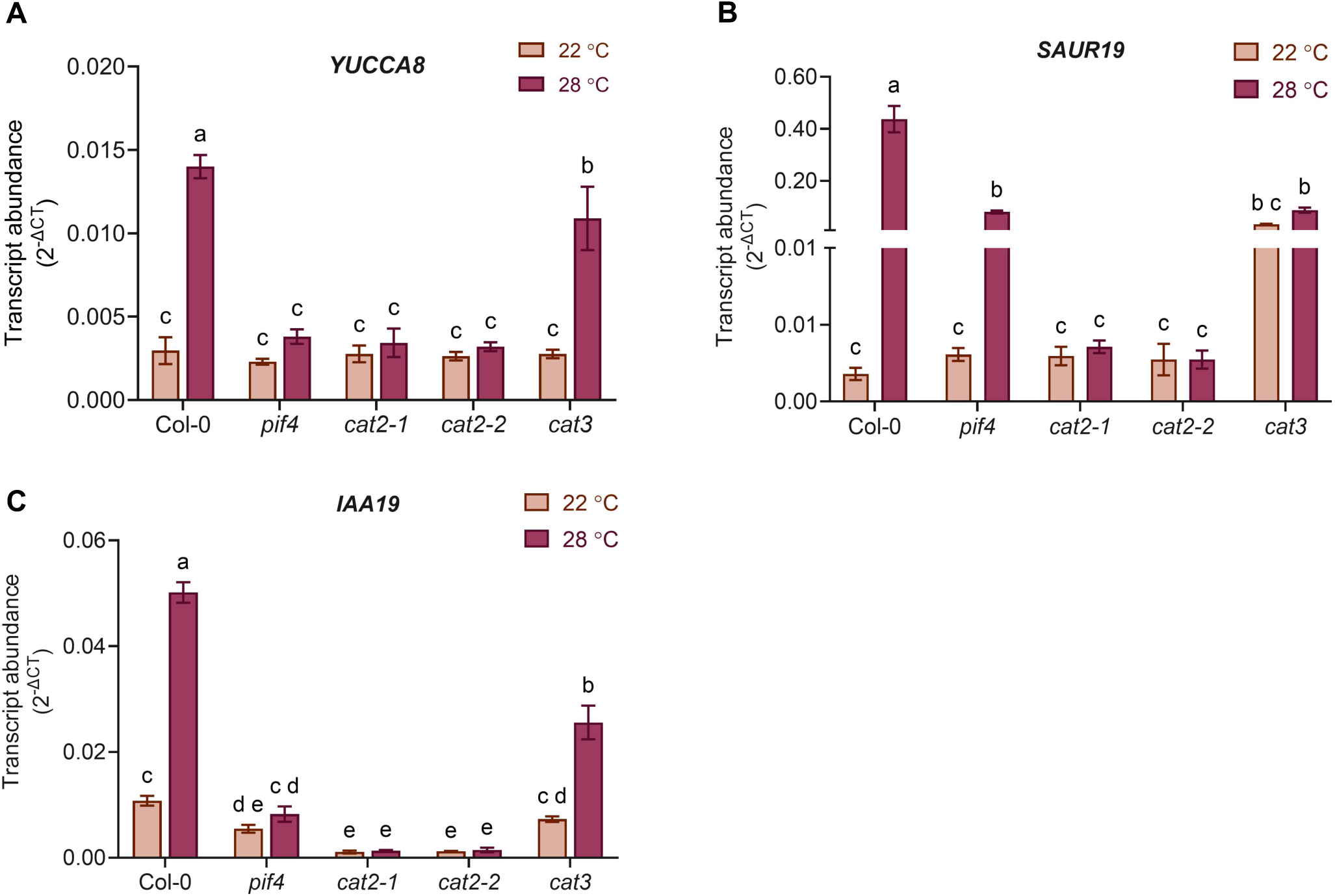
**Expression pattern of auxin biosynthesis and signaling genes in *cat* mutants along with Col-0 and *pif4* controls**. Relative transcript abundance (2_-ΔCT_) of auxin biosynthesis gene *YUCCA8* (A) and auxin signaling genes *SAUR19* (B) and *IAA19* (B), normalized to endogenous control *18S* in Col-0 and *pif4* along with *cat* mutants seedlings grown under control (22 °C) and high temperature (28 °C) conditions. Error bar represents SD (n=6). Different lowercase letters indicate statistically significant differences among the samples, as determined by two-way ANOVA followed by Tukey’s post hoc test (*P*<0.01).

